# Parallel and nonparallel genomic responses contribute to herbicide resistance in *Ipomoea purpurea*, a common agricultural weed

**DOI:** 10.1101/647164

**Authors:** Megan Van Etten, Kristin M. Lee, Shu-Mei Chang, Regina S. Baucom

## Abstract

The repeated evolution of herbicide resistance has been cited as an example of genetic parallelism, wherein separate species or genetic lineages utilize the same genetic solution in response to selection. However, most studies that investigate the genetic basis of herbicide resistance examine the potential for changes in the protein targeted by the herbicide rather than considering genome-wide changes. We used a population genomics screen and targeted exome re-sequencing to uncover the potential genetic basis of glyphosate resistance in the common morning glory, *Ipomoea purpurea*, and to determine if genetic parallelism underlies the repeated evolution of resistance across replicate resistant populations. We found no evidence for changes in 5-enolpyruvylshikimate-3-phosphate synthase (*EPSPS*), glyphosate’s target protein, that were associated with resistance, and instead identified five genomic regions that show evidence of selection. Within these regions, genes involved in herbicide detoxification--cytochrome P450s, ABC transporters, and glycosyltransferases--are enriched and exhibit signs of selective sweeps. One region under selection shows parallel changes across all assayed resistant populations whereas other regions exhibit signs of divergence. Thus, while it appears likely that the physiological mechanism of resistance in this species is likely the same among resistant populations, we find patterns of both similar and divergent selection across separate resistant populations at particular loci.

## Introduction

The evolution of pesticide resistance is a key example of rapid evolutionary change in response to strong, human-mediated selection [1]. Due to the widespread use of insecticides and herbicides in agriculture, multiple resistant pest populations often exist across the landscape [2–4]. These repeated examples of resistance allow for questions about the level at which parallel adaptation occurs [5–7]—*e.g.*, are parallel resistant phenotypes in separate lineages due to parallel changes at the developmental, physiological, or genetic level? Herbicide resistant weeds in particular provide remarkable examples of evolutionary parallelism, since the same nucleotide change can lead to resistance among separate lineages and even separate species [1,8,9]. Further, these examples of ‘extreme parallelism’ are often broadly considered as evidence of genomic constraint [7,10], which is the idea that parallel phenotypic evolution occurs because there are a finite number of genetic solutions to the same, often novel, environmental pressure.

Among herbicide resistant plants, the data that support the constraint hypothesis stems from sequence analysis of genes that are *a priori* known to produce the protein targeted by the herbicide (*i.e*., cases of target site resistance, TSR [9]) rather than genome-wide sequence surveys such as population genomics scans or genetic mapping studies. As a result, we understand very little about the potential for parallel genetic responses that may occur across the genome beyond the potential for changes within the (most often) single genes responsible for TSR. This is problematic as many weed species exhibit non-target-site resistance (NTSR) [11], which is caused by any physiological mechanism that is not due to TSR. NTSR can include a range of mechanisms, from herbicide detoxification to transport differences to vacuole sequestration [11]. Intriguingly, some weed species show multiple NTSR mechanisms within a single lineage [2,12,13], and even evidence of both TSR and NTSR [2,14]. Because there are relatively few examples underscoring the genetic basis of NTSR in herbicide resistant plants, it is currently unclear how ubiquitously cases of herbicide resistance support the idea of extreme genetic parallelism.

Previous research on the genetic basis of glyphosate resistance in crop weeds has focused largely on the potential for changes at the target site, the enzyme 5-enolpyruvylshikimate-3-phosphate synthase (*EPSPS*), which is a central component of the shikimate acid pathway in plants [15]. Conformational changes to the enzyme, due to mutations in the EPSPS locus, lead to target site resistance (TSR). There are also nontarget site resistance mechanisms responsible for glyphosate resistance in other weeds [11]; however, unlike the cases of resistance controlled by TSR, the genomic basis of NTSR to glyphosate has been characterized in very few species [16]. As a result, it is unknown if the same genetic basis underlies NTSR mechanisms across separate resistant populations. Thus, examining the genomic basis of resistance among replicated, resistant weed populations would provide an ideal study system to interrogate the hypothesis that genomic constraint underlies the parallel, repeated evolution of the resistance phenotype.

*Ipomoea purpurea* is a common agricultural weed that shows both within-and among-population variation in the level of resistance to glyphosate, the active ingredient in the widely used herbicide RoundUp: while some populations of this species across its range in the southeastern and Midwest United States exhibit high survival following herbicide application (high resistance), other populations exhibit low survival (high susceptibility) [4]. The pattern of resistance across populations suggests that resistance has evolved repeatedly, with highly susceptible populations interdigitated among resistant populations [4], and no evidence of isolation-by-distance across populations, as would be expected in the simple scenario wherein resistance evolved once and moved across the landscape *via* gene flow [4]. We have recently shown that neutral genetic diversity across these populations is negatively related to the level of resistance [17] and that additive genetic variation underlying resistance to glyphosate in *I. purpurea* responds to selection *via* the herbicide [18,19]. Additionally, there is evidence of a fitness cost associated with glyphosate resistance in the form of lower seed germination and smaller plant size [20]. Intriguingly, the resistant populations appear to vary in the expression of this cost -- some highly resistant populations exhibit low germination and others exhibit smaller size, on average, than susceptible populations [20]. These data suggest that perhaps the genetic basis of resistance, or the physiological mechanism underlying resistance, differs among resistant populations. However, the genetic basis of resistance across any population of this species is currently unknown.

Our overarching goal is to determine if the same genetic basis is responsible for glyphosate resistance across separate populations of *I. purpurea* sampled from agricultural fields with a history of glyphosate exposure. We first evaluate the potential for sequence changes in the EPSPS locus and find there are no changes that correlate with resistance, providing evidence that target site resistance is not responsible for the resistance phenotype across populations. We then perform a population genomics screen to identify loci that exhibit signs of selection--thus putatively responsible for the resistance phenotype--and to determine if patterns of relatedness between resistant populations suggest a single or multiple origins of resistance. We follow up on this screen with exome resequencing of candidate resistance loci, and determine if populations share a similar haplotype structure, which would suggest that a similar genetic basis was responsible for resistance across the landscape. We find regions of the genome that show evidence of selection across resistant populations to contain genes responsible for herbicide detoxification. Additionally, patterns of haplotype sharing among populations suggests both parallel and nonparallel genomic responses underlie resistance among populations. Overall, our results suggest that evolutionary constraints can underlie herbicide adaptation, but that patterns of selection across the genome indicate the potential for both parallel and divergent responses.

## Results

### No evidence for changes in glyphosate target enzyme (*EPSPS*)

We sequenced two copies of *EPSPS* (copy A and B) from geographically separate populations of *I. purpurea* to determine if glyphosate resistance is due to a target-site resistance mechanism in this species as identified in other resistant species [21]. Individuals used for sequencing were sampled as seed from six highly resistant (R) (N=20, average survival at 1.7kg a.i./ha: 84%) and five susceptible (S) populations (N=25, average survival at 1.7kg a.i./ha: 26%; S1 Table) [4]. We found 14 (copy A) and 22 (copy B) variable sites across all populations but no copy exhibited SNPs in the region previously shown to cause resistance in other weed species (S1 Fig). Additionally, resistant and susceptible populations did not significantly differ in allele frequencies for any of these SNPs (copy A: chi-squared test, χ^2^ range 0.02-0.33, min p-value = 0.57; copy B: chi-squared test, χ^2^ range 0.00-0.18, min p-value = 0.67; S1 Table) nor were any significantly correlated with resistance level (copy A: Pearson’s correlation, coefficient range 0.25-0.69, min p-value = 0.12; copy B: Pearson’s correlation, coefficient range 0.15-0.72, min p-value = 0.17; S1 Table).

### Population structure suggests independent evolution of resistance

We next examined measures of genetic relatedness to determine if separate resistant populations showed a pattern of high similarity, which would suggest that resistance alleles were shared between populations due to gene flow or from a common lineage. To do so, we used a modified RAD-seq approach (nextRAD) and genotyped 10 individuals sampled as seed from each of four resistant populations and four susceptible populations (average survival at 1.7kg a.i./ha: 89% and 16%, respectively [4]; Fig 1A; Table 1). This resulted in 8,210 high-quality, variable SNP loci from 80 individuals. Population genetics parameters of the RADSeq SNPs, including expected and observed heterozygosity across populations are presented in the S2 Table. A neighbor joining tree calculated from pairwise relatedness showed that resistant populations did not cluster into a single group and are instead interspersed with the susceptible populations (S2 Fig). Additionally, a principal coordinates analysis (PCoA) using allele frequencies (Fig 1B) did not separate the populations into distinct resistant and susceptible groups, and a genetic structure analyses showed that resistant and susceptible populations did not segregate into two separate genetic clusters as would be expected if all resistant populations derived from the same initial population (S3 Fig).

**Table 1.** Population information for each population used in the study. Pop number = population number as used in other studies resulting from this seed collection, Resistance type = classification of resistance in the population R >0.5 prop. survival S<0.5 prop. Survival, Pop Abbrev. = abbreviation for each population as used in other studies, State = state where seeds were collected, Proportion survival at 1.7 = proportion of individuals that survived a spray rate of 1.7 kg/ha of glyphosate based on Kuester et al 2014, Latitude and Longitude = location where seeds were collected, Used for = abbreviation for which part of the study each population was used for E=EPSPS sequencing P=population genetics.

| Pop number | Resistance type | Pop Abbrev. | State | Proportion survival at 1.7 | Latitude | Longitude | Used for |
| --- | --- | --- | --- | --- | --- | --- | --- |
| 42 | S | SH4 | VA | 0.1 | 38.373523 | -78.662516 | E,P |
| 4 | S | CR | NC | 0.21 | 34.556672 | -79.125602 | E |
| 36 | S | IN12 | IN | 0.25 | 40.565608 | -85.503826 | E |
| 17 | S | MA1 | SC | 0.25 | 34.159155 | -79.272908 | E |
| 23 | S | SN | TN | 0.5 | 35.067905 | -86.62955 | E |
| 19 | R | MC | NC | 0.67 | 34.508193 | -78.70899 | E |
| 5 | R | CL1 | SC | 0.73 | 33.859875 | -79.909072 | E |
| 43 | R | VA2 | VA | 0.82 | 36.886448 | -78.553156 | E |
| 32 | R | WG | TN | 0.83 | 35.099356 | -86.225509 | E,P |
| 1 | R | BI | TN | 1 | 35.775237 | -85.903419 | E,P |
| 10 | R | DW | NC | 1 | 34.983161 | -78.039309 | E,P |
| 48 | S | RB | TN | 0.18 | 35.31653 | -87.35373 | P |
| 14 | S | HA | NC | 0.15 | 35.424763 | -77.917121 | P |
| 12 | S | FL | SC | 0.20 | 34.145812 | -79.865313 | P |
| 51 | R | SPC | TN | 0.71 | 35.533413 | -85.951902 | P |

**Fig. 1.**
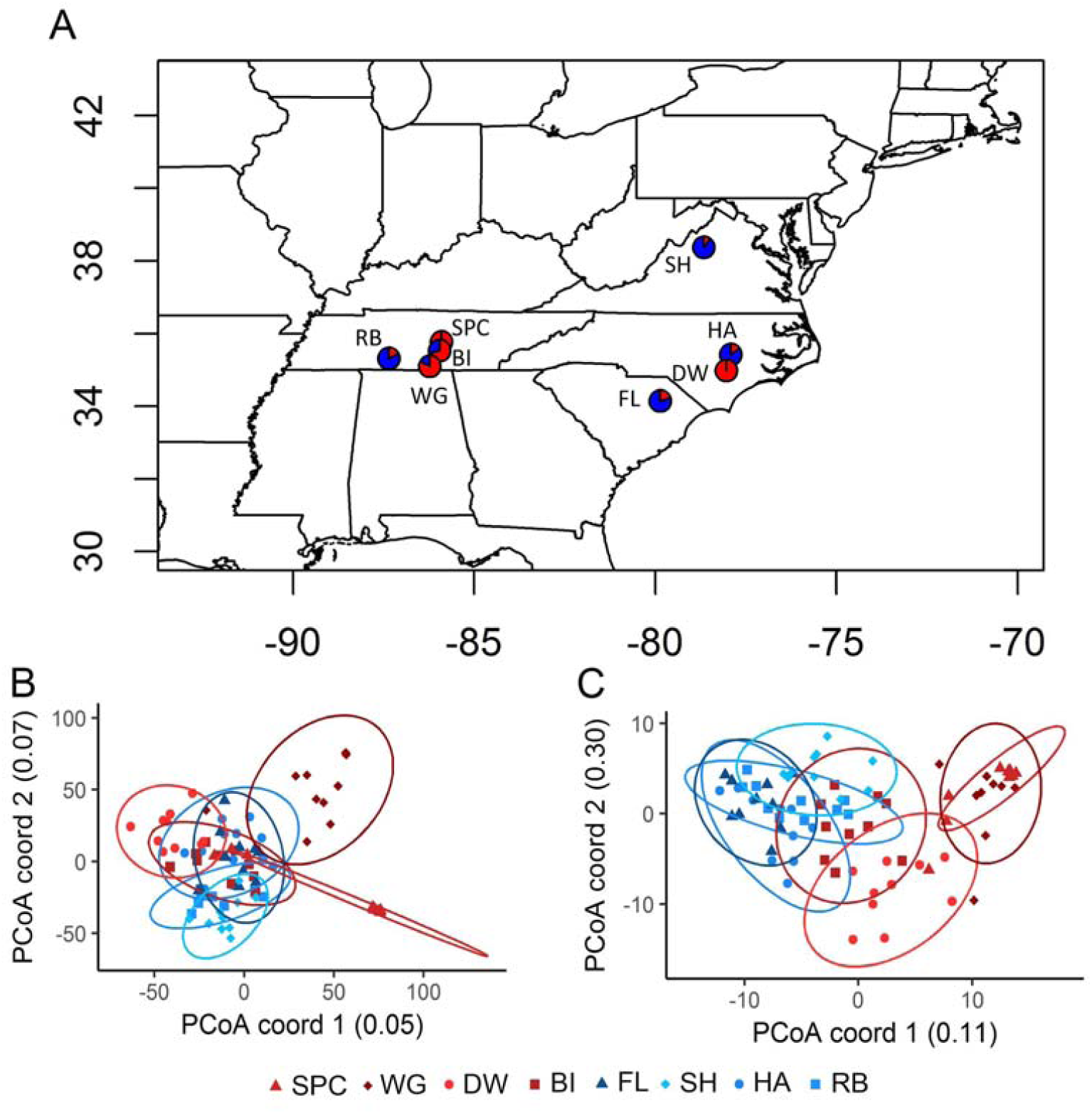
Population locations and relationships among *I. purpurea* samples. (A) Populations were sampled from locations in the southeast and ranged from 10% to 100% survival following glyphosate application (proportion of individuals that survived glyphosate treatment shown for each population, red=survived, blue=died). Individuals from resistant populations (>50% survival after treatment; red colored symbols) do not group together in a PCoA analysis (B) when using all of the RAD-seq SNP loci. Allele frequencies of outlier loci are presented in (C). Populations indicated in blue are highly susceptible whereas populations in red are resistant to glyphosate.

### Genome-wide scan indicates loci associated with resistance

We next performed a genome-wide outlier screen to identify loci exhibiting signs of selection and thus potentially involved in glyphosate resistance in *I. purpurea.* We used two programs (BayeScan and bayenv2) to do so. BayeScan identified 42 loci that were outliers while bayenv2 identified 83 loci whose allele frequencies were correlated with the level of resistance (Dataset S1). Using GO assignments (Dataset S1), we found that the top three biological processes for the resistance outlier loci were proteolysis, protein phosphorylation, and regulation of transcription. Of special note, we identified a glycosyltransferase among the outlier loci, which are genes shown to be involved in herbicide detoxification in other species [12,22,23].

The identified resistance outliers showed twice the level of differentiation among the resistant populations (mean pairwise F_ST_s of outliers = 0.327, 95% CI = 0.293-0.362) compared to the level of differentiation among susceptible populations (mean pairwise F_ST_s of outliers = 0.180, 95% CI = 0.146-0.216). This contrasted with genome-wide patterns of F_ST_ (*i.e.* pairwise F_ST_ across all loci: resistant populations F_ST_ = 0.198 (0.192-0.203), susceptible populations F_ST_ = 0.133 (0.128-0.137)). Further, the pattern was the same for outliers regardless of whether they were identified by BayeScan or bayenv2. This increased differentiation of outlier loci among resistant populations could be a result of drift, or could indicate that a different genetic basis underlies resistance across populations. Two resistant populations from central Tennessee (SPC and WG) exhibited significant overlap in allele frequencies of outlier loci (Fig 1C), suggesting a similar response to selection between these two populations. On the other hand, the allele frequencies of outliers from BI, another highly resistant population from TN, clustered between the susceptible and other resistant populations whereas individuals from DW, a resistant population from North Carolina, exhibited some overlap with BI (Fig 1C).

To insure that our resistance outliers from the RADseq analysis were associated with resistance rather than an environment that might co-vary with the level of resistance, we examined three other likely environmental variables in a separate bayenv2 analysis: minimum temperature of the coldest month, precipitation of the driest month, and elevation. We chose these specific climatic variables as other herbicide resistance studies have identified the influence of temperature and precipitation on the expression of resistance within a population [24–27]. While this tactic identified loci that were associated with environmental variables, very few of these loci overlapped with our identified resistance loci, indicating that the loci that are associated with resistance are not likely the result of selection by other environmental influences (S4 Fig).

### Exome re-sequencing identifies genomic regions associated with resistance

We next performed target-capture re-sequencing of the genes located near (or containing) outlier SNPs identified by the population genomics screen. Using both a *de novo* genome and transcriptome assembly [28] (S3 Table), we designed probes to sequence the following: exons from predicted genes near outlier SNPs (171 genes), genes from a match of the outlier SNPs to the transcriptome (30 genes), the *EPSPS* genes (2), previously reported differentially expressed genes associated with resistance [28] (19 genes), and 214 randomly chosen transcriptome sequences to serve as a control (Dataset S1). We made target-enriched libraries for 5 individuals in each of the 8 populations (Fig 1A), which were then sequenced on an Illumina Hi-Seq 2000. Following sequencing, filtering, and contig assembly (see Methods) we ran outlier tests to identify SNPs exhibiting signs of selection. Of this set, BayeScan identified 104 SNP outliers while bayenv2 identified 231 SNP outliers, 98 of which were shared between programs (Dataset S1). The majority of outliers were from the probes designed from the population genomics RAD-seq outliers (52%), followed by the non-probe contigs (*i.e.* off-target sequences; 37%), and a few from the control probes (11%). The majority of the outliers from control probes (17/20) fell within genomic regions that we found to be enriched with outliers (*see below*).

We aligned the re-sequenced contigs onto the assembled genome of a close relative, *I. nil* [29], and identified five genomic locations that were enriched for outliers (Fig. 2A), with 149 (71%) of the outlier SNPs falling within these regions. The five regions ranged from 276 KB to 4 MB in size and together contained 945 predicted genes (based on *I. nil* gene annotations; Dataset S1). Some of the five regions contained outliers identified by both bayenv2 and BayeScan while others regions had outliers primarily identified by bayenv2 (% of outlier SNPs identified by both programs, chromosome 1: 6%; chromosome 6: 72%; chromosome 10: 100%;chromosome 13: 60%; chromosome 15: 36%). The outlier enriched regions were not located near or within the centromere for any chromosome (centromere indicated by thick vertical line on the x-axis, Fig 2A).

**Fig 2.**
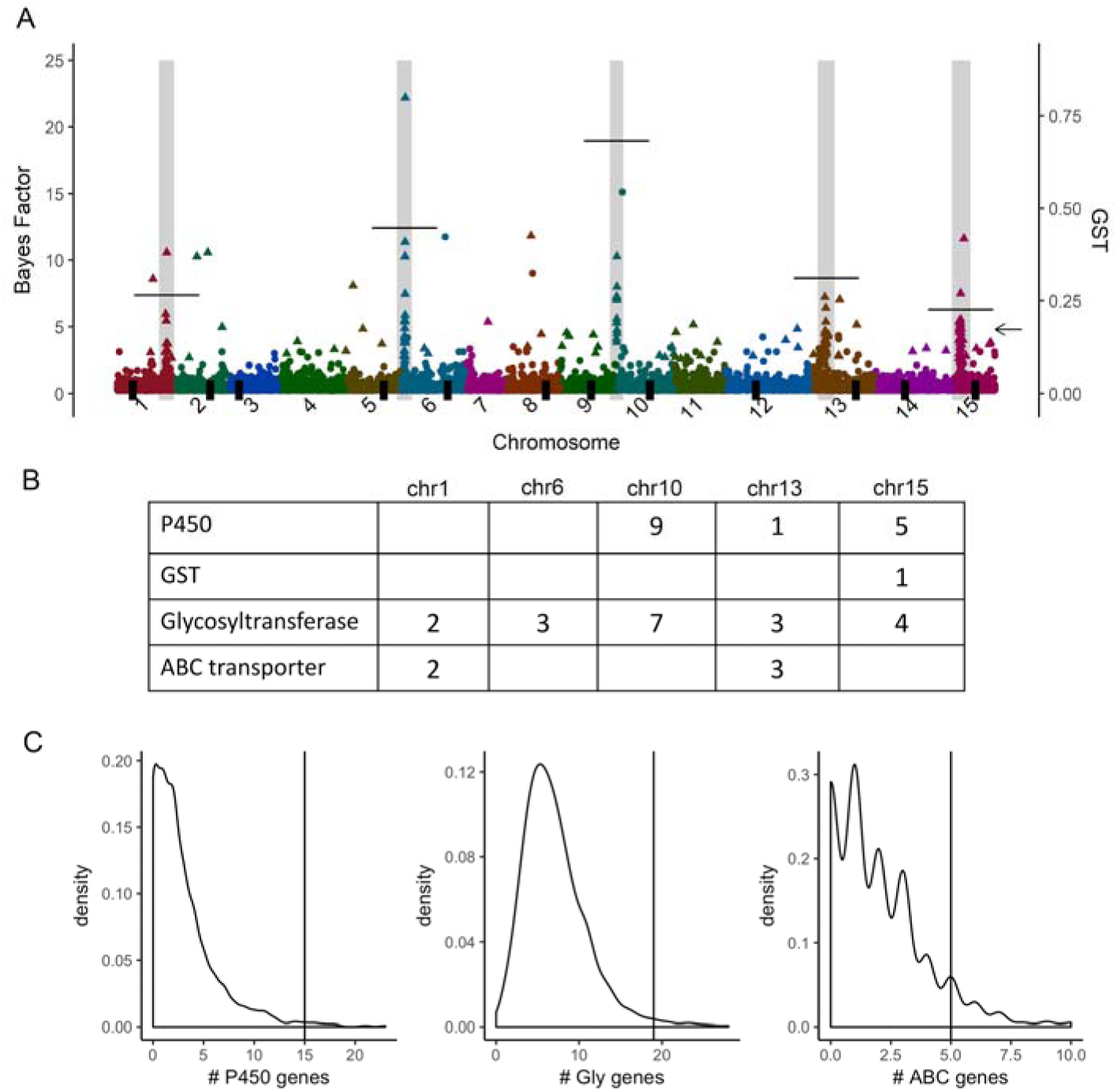
Regions of the *I. purpurea* genome enriched with outlier loci. (A) Aligning the denovo contigs to the *I. nil* genome shows 5 regions enriched for outliers (regions in grey; symbol colors denote chromosomes; symbol shape denotes significance). The majority of the outliers (71%) fall within the five regions. Significant outliers, noted with triangles, exhibited the most extreme 1% Bayes Factors and the 5% most extreme Spearman correlation coefficients (left y-axis). The average GST (right y-axis) was calculated per enriched region and is indicated by a thin horizontal line for each outlier enriched region (arrow indicates average GST value over all SNPs). The position of each chromosome’s centromere is indicated by a thick black vertical line on the x-axis. (B) The five outlier-containing regions (chr1-chr15) had multiple copies of several genes potentially involved in non-target site resistance (numbers indicate the number of genes that fall into each category, P450 = cytochrome P450, GST = glutathione s-transferase, Glycosyltransferase = glycosyltransferase, ABC transporter = ABC transporter). (C) Resampling the *I. nil* genome 1000 times to generate an empirical distribution of gene copy number of each type of gene indicates that the outlier enriched regions contain more of the potential herbicide detoxification genes of interest than expected due to chance. Thin horizontal line indicates overall number of each type of gene found within the outlier-enriched regions, which was greater than expected from the empirical distribution for the cytochrome P450, glycosyltransferase, and ABC transporter genes (P < 0.001).

We identified multiple genes within the outlier enriched regions from four gene families of interest—the cytochrome P450s, ABC transporters, glycosyltransferases, and glutathione S-transferases (GST)—which are gene families hypothesized to be involved in non-target site resistance *via* herbicide detoxification (Fig 2B). Resampling 1000 times identified a significant over-representation of glycosyltransferase (P = 0.01), ABC transporter (P = 0.05), and cytochrome P450 (P = 0.01) genes within the five enriched regions (Fig 2C), suggesting that these loci are potentially responsible for resistance in *I. purpurea* and were not identified solely due to their high copy number in plant genomes. In comparison, outlier SNPs that did not fall into the five outlier enriched regions (29% of SNPs) were less likely to be near genes from these four families (S5A-D Fig).

As expected based on the Bayescan results, the regions of each of the five chromosomes enriched with outliers exhibited high genetic differentiation between resistant and susceptible populations (average across genome is indicated by the arrow on Fig 2A; measured as G_ST_, which is F_ST_ generalized to multiple alleles). Although all regions showed an average G_ST_ > 0.20, the enriched region on chromosome 10, spanning ∼0.28MB, displayed the highest G_ST_ (chr 10 enriched region avg+SD: 0.64+0.12, R vs S populations). Within this region, we found higher nucleotide diversity among susceptible compared to resistant individuals (π_S_/π_R_ = 2.04; a ratio more extreme than that found across 95% of the genome-wide SNP windows, Fig 3A; S6 Fig). In comparison, across other outlier enriched regions, nucleotide diversity was higher among resistant compared to susceptible individuals, but the difference between resistant and susceptible individuals exceeded the background genome-wide ratio only within the enriched region on chromosome 1 (Fig 3A).

**Fig 3.**
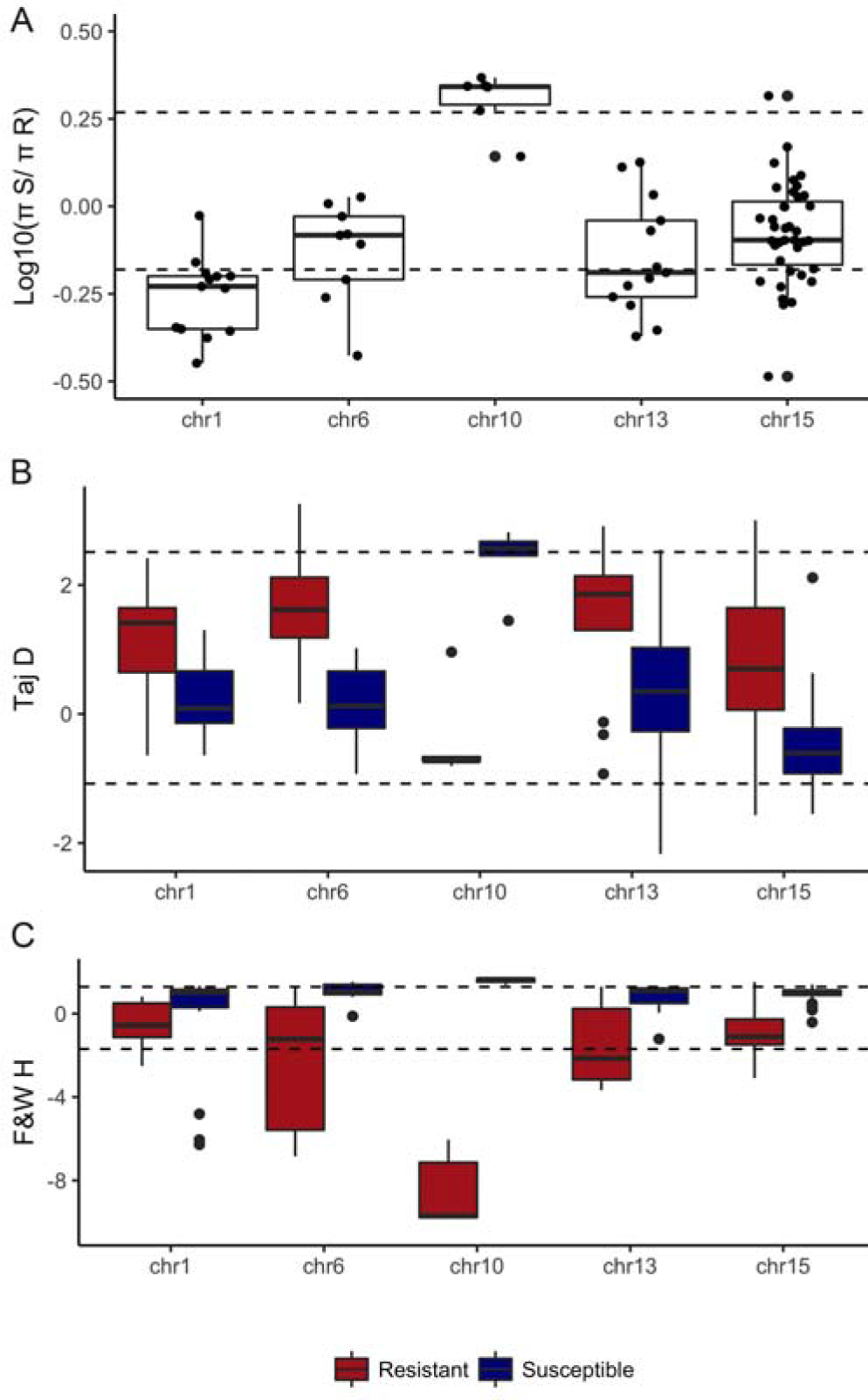
Resistant individuals exhibit evidence of selective sweeps in outlier-enriched regions of genome. (A) Nucleotide diversity (shown here as log10 piS/piR) is decreased in resistant individuals within the chr10 region compared to susceptible individuals, and (B) values of Tajima’s D and (C) Fay and Wu’s H across outlier enriched regions both suggest marks of positive selection in the chromosome 10 outlier enriched region, with some indication for positive selection in the outlier enriched region of chromosome 13. Dashed lines show the 95% most extreme genome-wide values for each metric.

The outlier enriched region on chromosome 10 likewise exhibited evidence of selection based on estimates of both Tajima’s D (Fig 2B) and Fay and Wu’s H (Fig 2C). Tajima’s D, which is sensitive to a lack of low-frequency variants [30], exhibited a negative value among resistant individuals, although the most extreme values within this region ranged from −0.64 to −0.81 and did not exceed the 95% most extreme genome-wide values (Fig 2B). In comparison, Fay and Wu’s H, which is sensitive to excess high-frequency derived variants compared to neutral expectations [31], was significantly more negative than the genome-wide value among resistant individuals (−8.55; Fig 2C). Interestingly, values of Tajima’s D and Fay and Wu’s H were typically positive and either greater than 2 (2.37, avg Tajima’s D in region) or approaching 2 (1.59, avg Fu and Way’s H in region) among susceptible individuals, suggesting a pattern of balancing selection within susceptible populations. The difference in both Tajima’s D and Fu and Way’s H between resistant and susceptible individuals within two 25 SNP windows (positions 381983679 - 382012084) were more extreme than that found across 99% of the genome-wide SNP windows, potentially narrowing in to a ∼28 kb region of strong selection within the outlier enriched region of chromosome 10. Finally, the enriched region on chromosome 13 exhibited negative values of Fu and Way’s H among resistant individuals (−1.58, avg Fu and Way’s H within region), with the most extreme negative values ranging from −2.15 to −3.68 over a contiguous region of 1.49MB.

Given signs of positive selection on the outlier enriched regions of chromosome 10 and (to a lesser extent) chromosome 13, we examined the genes found within these two regions in greater detail. Within the outlier-enriched region of chromosome 10, we identified 7 glycosyltransferase and 9 cytochrome P450 genes, with the 7 glycosyltransferase genes found tandemly repeated within a span of 42 kb (Fig 4A). Seventeen non-synonymous SNPs were present across four of the glycosyltransferase genes (asterisks in Fig 4A). Within an 811 bp segment of the conserved domain one of the glycosyltransferases, we identified a cluster of seven non-synonymous SNPs with very low π values in resistant compared to susceptible individuals (conserved domain average π_R_ = 0.18; π_S_ = 0.43). None of the non-synonymous SNPs within this region were fixed within the resistant populations, but were very close to fixation (allele 1, resistant freq = 0.1, susceptible freq = 0.7; allele 2, resistant freq = 0.9, susceptible freq = 0.3). Within the outlier-enriched region of chromosome 13, we identified a cytochrome P450 gene with 6 non-synonymous SNPs (shown with asterisks in Fig 4B), and a shared haplotype among three of the four resistant populations (Fig 4B).

**Fig 4.**
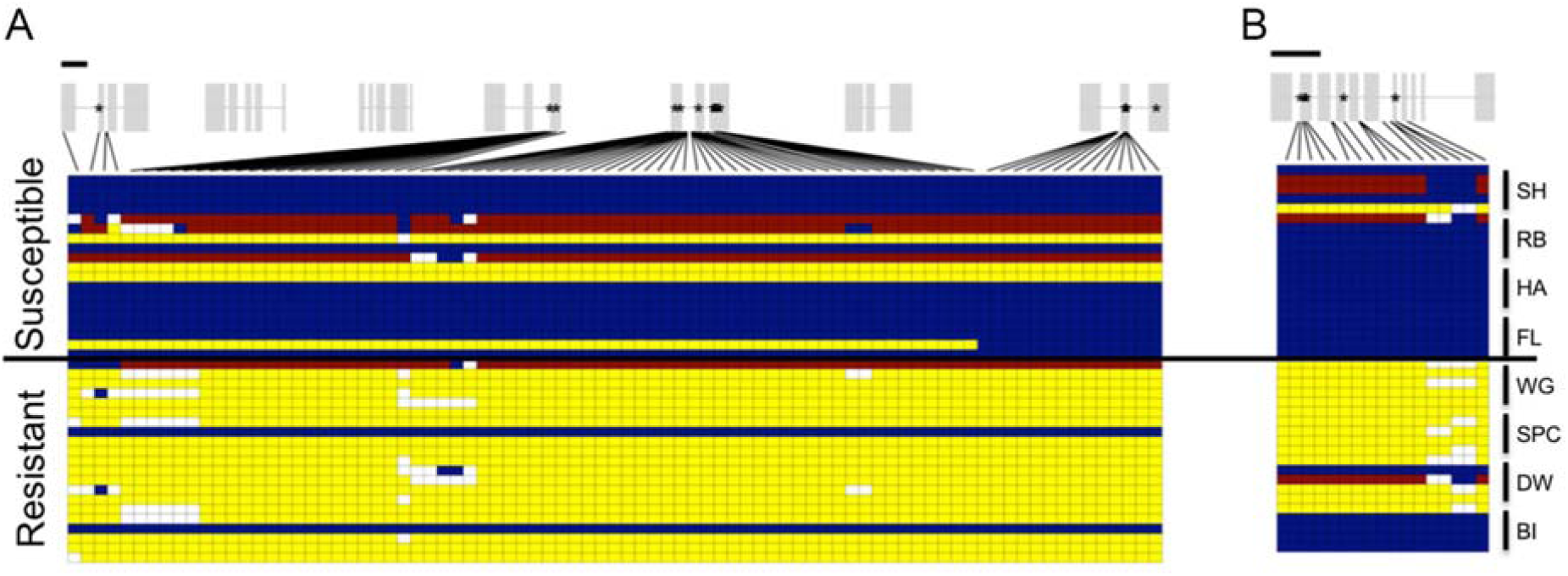
Signs of selection across conserved haplotypes of herbicide detoxification genes. Haplotypes are shown for each individual for the (A) seven duplicated glycosyltransferase genes on chromosome 10 (exons above in grey), and (B) an ABC transporter gene on chromosome 13. Blue and yellow indicate homozygotes, red indicates heterozygotes, white in missing data; stars indicate a non-synonymous change at that location. Black bar above gene models indicates 1kb.

We likewise examined patterns of linkage disequilibrium across the outlier enriched regions of each of the five chromosomes, since linkage between SNPs would provide another line of evidence for a potential selective sweep indicating a response to selection. Additionally, we calculated linkage disequilibrium (LD) along the chromosome (for chrs 1, 6, 10, 13 and 15) to determine an expected background amount of linkage between SNPs and thus an idea of the efficacy of our RADseq followed by exome-resequencing approach for identifying the genetic basis of resistance among populations. Across each chromosome, we found the average r^2^ values (the correlation coefficient between each SNP pair as our estimate of LD) to be low, ranging from 0.032-0.036 (S4 Table). Due to the granular nature of the data, we did not estimate linkage decay, but did examine the potential for linkage within 10 kb windows on average. These values were greater than the background LD, but still less than 0.1 (range 0.038-0.078, S4 Table). In comparison to values of linkage across the entire chromosome, we found evidence of stronger linkage among SNPs within the outlier-enriched regions of chromosomes 1, 6, 10, 13 and 15 (range of average r^2^, 0.12-0.23). Notably, the chromosome 10 outlier-enriched region exhibited the highest r^2^ value (0.234, S5 Table). Because the outlier-enriched regions varied in length, thus complicating the comparison of LD between them, we qualitatively examined the length around each outlier enriched region with elevated LD, or r^2^ values that were > 0.25. We found that each outlier enriched region exhibited r^2^ > 0.25 across relatively large sequence lengths, which ranged from 84 kb to 3 MB across chromosomes (S5 Table).

### Haplotype structure

A goal of the present work was to determine if separate populations have responded in parallel at the genomic level to selection *via* herbicide application. We performed a visual examination of the haplotype structure among outlier-enriched regions in more depth with the idea that a similar haplotype among separate resistant populations would point to a shared genomic basis underlying at least some of the loci indicated in herbicide resistance and another indication of selection on those loci. We used hierarchical clustering for this examination of haplotype structure. Using each sequenced contig from the outlier-enriched regions (Chrs 1, 6, 10, 13, and 15), we assigned individuals to one of two groups based on genetic distance—either the group that contained the majority of susceptible individuals from highly susceptible populations (hereafter the ‘S’ group) or the other group (hereafter the ‘R’ group). We found a high proportion of resistant individuals (>75%) across all four resistant populations (SPC, WG, DW, and BI) in the chromosome 10 outlier enriched region (Fig 5A), meaning that the majority of resistant individuals from these populations shared high levels of genetic similarity in this region. Likewise, a high proportion of resistant individuals exhibited high genetic similarity in the outlier enriched region on chromosome 6, but only in three of the four resistant populations (SPC, WG, and DW). In contrast, the enriched regions on chromosomes 1, 12 and 15 exhibited high proportions of resistant individuals for SPC and WG, but not BI and DW (Fig 5A).

**Fig 5.**
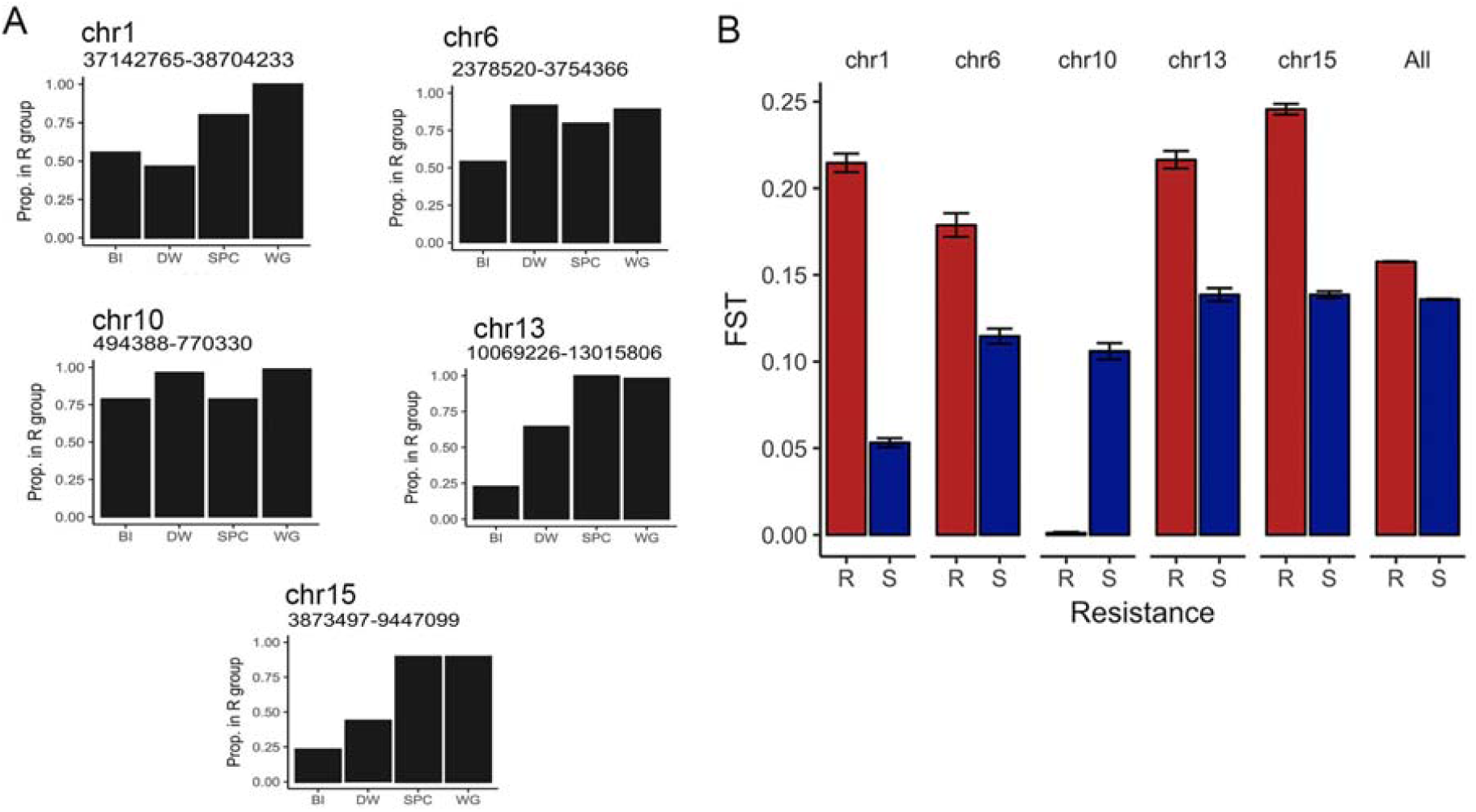
Genetic similarity of haplotypes among resistant populations. (A) The proportion of each population that exhibited the resistant haplotype are shown for each population. Pairwise genetic distance between each individual was calculated using all SNPs from each *I. purpurea* contig from the outlier-enriched regions (length of contig used shown for each chromosome), and multidimensional scaling was used to reduce the resultant genetic distance matrix to two dimensions. Populations were then hierarchically clustered into two groups, with the group containing less than half of the individuals from the susceptible populations considered the ‘resistant’ group. (B) The average pairwise genetic differentiation for resistant (red) and susceptible (blue) populations. Pairwise FST values were calculated separately for resistant and susceptible populations using contigs from each outlier enriched region of each chromosome.

Additionally, we examined patterns of pairwise genetic differentiation among resistant and susceptible populations of the outlier-enriched regions of each chromosome, with the general expectation that a higher pairwise F_ST_ between resistant populations, compared to susceptible populations, might indicate lack of gene flow and/or greater genetic differences between resistant populations within these regions. We calculated pairwise F_ST_ estimates [32] among the resistant populations and the susceptible populations separately for each SNP, and then compared the average pairwise F_ST_ of the resistant populations versus the susceptible populations within the 5 outlier enriched regions. Across chromosomes 1, 6, 13, and 15, we found higher pairwise F_ST_ among resistant populations compared to susceptible populations, indicating that resistant populations were more differentiated in these regions. On chromosome 10, in comparison, we found no evidence of genetic differentiation among resistant populations, suggesting either strong selection on young standing genetic variation within this region among populations, or the potential that gene flow has recently occurred between them followed by subsequent recombination (Fig 5B).

### Formal test of convergence

Given multiple lines of evidence suggesting the region on chromosome 10 has responded in parallel across the examined resistant populations (*i.e*., an outlier enriched region with high differentiation between resistant and susceptible populations, a similar haplotype among resistant populations, marks of selection based on nucleotide diversity, Tajima’s D, and Fu and Way’s H, and evidence for linkage between markers within the enriched region), we next performed tests to examine the nature of convergence within this region. More specifically, we sought to determine the most likely model for genomic convergence by determining whether potential selected alleles within the region on chromosome 10 exhibited multiple independent origins, were spread among populations *via* gene flow, or were shared among populations due to ancestral standing variation. To do so, we applied the inference method of Lee and Coop (2017), which builds on coalescent theory to show how shared hitchhiking events influence the covariance structure of allele frequencies between populations at loci near the selected site. Although our screens indicated multiple regions of the genome under selection, in this work we focus formal tests of convergence only on the enriched region of chromosome 10 given the evidence for a high proportion of individuals exhibiting the same haplotype. This pattern is suggestive of a selective sweep that was shared among resistant populations, and one that was due to selection on young standing, shared genetic variation, or due to migration between populations.

We applied the inference method using 2248 SNPs from the ∼276 KB region of the contig encompassing the outlier enriched regions on chromosome 10 to identify the locus under selection and to distinguish the most likely model of adaptation (independent, *de novo* mutations, migration, or selection on standing ancestral variation). From this analysis we find the migration and standing variation models to show similarly high log-likelihood ratios (Fig 6A). All three models peak at position 381,993,922 (based on the *I. nil* genome), indicating the most likely selected site. Notably, this position is within the two SNP windows that exhibited signs of selection from estimates of Tajima’s D and Fu and Way’s H (Fig 3). Further examination of the standing variant model at this position shows the parameters that result in the highest likelihood are very low standing allele frequency (g = 10^-6) and very high selection (s = 1), with the amount of time that the beneficial allele has been standing in the populations prior to selection, or t, estimated to be 5 generations (Fig 6B, C). This standing time is much smaller than the population split times (289K generations ago), so we assume migration in the model and the five generations are interpreted as the time between gene flow between populations and the onset of selection. We ran the model with a denser grid of t (0-10 generations) and found that the likelihood value was highest when t was equal to 0, indicating that the beneficial allele was immediately advantageous after introgressing and began sweeping rapidly within populations. In comparison, for the migration model, the parameters that result in the highest likelihood are a migration rate of 1 and high selection (s = 0.65). Overall, our analyses of this region strongly supports a model where gene flow introduced the beneficial allele(s) into populations, which then began sweeping quickly and immediately. A rapid sweep like that proposed here would not allow for recombination to break down the haplotype introgressing along with the selected allele. This fits our expectations from the haplotype patterns above since there is high similarity between resistant populations over long stretches of this region.

**Fig 6.**
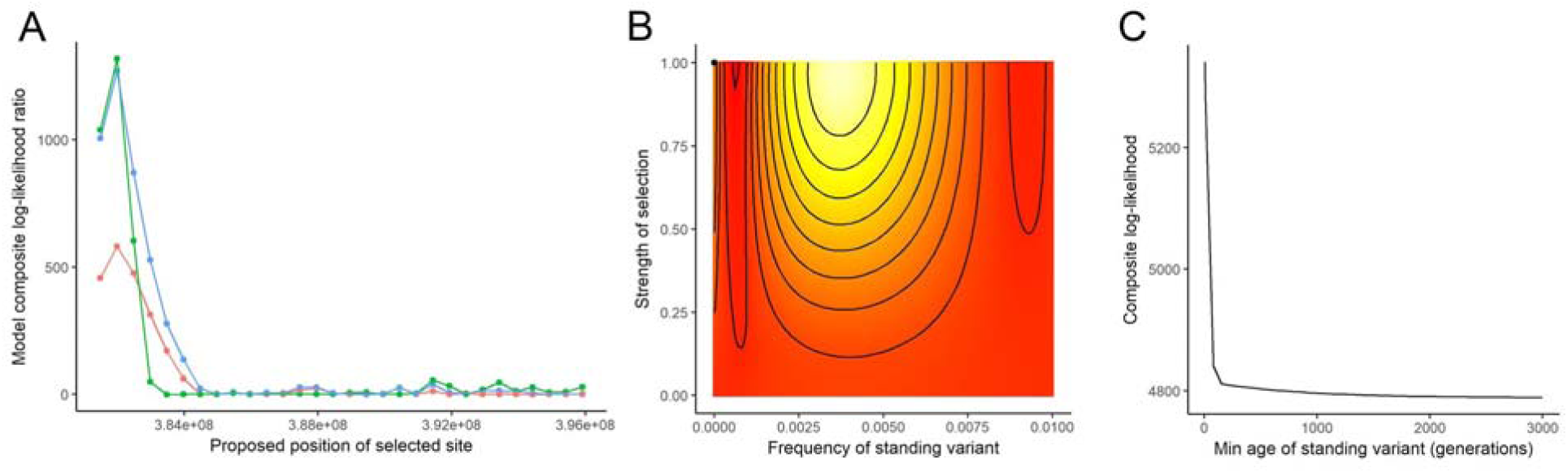
Test of convergence. (A) Likelihood ratio of the following models relative to a neutral model with no selection: standing variant model (blue), migration (green) or independent mutation (red). (B) Likelihood surface for minimum frequency of the standing variant and the strength of selection holding the age of the standing variant constant; the point indicates the highest likelihood. (C) Likelihood surface for the minimum age of standing variant maximizing over the other parameters.

## Discussion

In this work, we examined the evolution of glyphosate resistance across geographically separate populations of the common morning glory, *Ipomoea purpurea*. We set out to identify candidate loci involved in glyphosate resistance in this species and to determine if the pattern of selection on putative resistance loci was similar across highly resistant populations, which would indicate that populations responded in parallel to herbicide selection. Our results provide evidence that adaptation to glyphosate in *I. purpurea* is not due to a single gene, target-site resistance mechanism (TSR) as there are no nucleotide sequence differences in the target locus, *EPSPS*, that correlate with resistance. We found instead that at least five regions of the genome show evidence of selection and that these regions are significantly enriched for genes involved in herbicide detoxification. Further, we found evidence for a shared pattern of strong selection on one region of the genome among the four highly resistant populations (chromosome 10) whereas other regions under selection exhibited divergence between the resistant populations. These findings suggest that resistance in this species is due to a non-target genetic mechanism (NTSR), components of which exhibit signs of both parallel and non-parallel responses to selection among populations.

### Genetic basis of glyphosate resistance in *I. purpurea*

*Ipomoea purpurea* is a noxious crop weed found in disturbed agricultural sites in the Southeastern and Midwest US. Our previous work examining the level of resistance among 47 populations showed that resistance appeared on the landscape in a mosaic fashion, with highly resistant populations interdigitated among highly susceptible populations. This phenotypic pattern suggested resistance was independently evolving across populations [4]. Coalescent modelling using SSR marker variation supported a scenario of migration among populations prior to onset of glyphosate use (before 1974, when glyphosate was released commercially), rather than a scenario of migration *after* the introduction of the herbicide [4]. We thus hypothesized that resistance independently evolved among populations, and was most likely due to selection on standing and shared genetic variation [4]. However, we also found genetic differentiation among populations to be low (F_ST_ = 0.127; [4]), and a more recent fine-scale analyses of their connectivity showed that although the majority of individuals were sired from within populations, three of the resistant populations included in this work (WG, SPC, and BI) shared recent migrants [33]. These findings support the idea that migration between populations could allow for the sharing of resistance alleles. Both of these scenarios--migration prior to the widespread use of the herbicide, or very recent migration--suggest that resistance is likely to be controlled by the same genetic basis across populations. Intriguingly however, we also previously showed that fitness costs were different among resistant populations, suggesting that the genetic basis of resistance could potentially be different [20]. Thus, we used a sequencing approach across highly resistant but broadly separated populations to investigate the genetic basis of resistance and to determine if patterns of selection and haplotype sharing indicated that the same genomic features were responding to herbicide selection among populations.

We found no evidence supporting target site resistance in *I. purpurea*--there were no variants within the *EPSPS* locus associated with resistance, and we found no evidence for selection on copies of *EPSPS* from the exome resequencing data. Using both RNAseq and rtPCR, we previously showed that transcripts of EPSPS are not overexpressed in *I. purpurea* [28], providing evidence that resistance is not related to EPSPS overexpression, as has been shown in a variety of resistant species [34–38].

Given the lack of structural or expression-related changes to the target-site locus, *EPSPS*, we combined a population genomics screen and exome resequencing to identify potential candidate loci underlying resistance. This strategy identified five candidate regions of the genome that were enriched with loci exhibiting signs of selection. The pattern of genomic differentiation within these five regions was greater than that of genome-wide, background differentiation--suggesting a response to herbicide selection. None of these regions were physically located near the centromere, which has been shown in other species to be areas of reduced recombination and thus high differentiation [39–42]. We identified the strongest evidence for positive selection associated with resistance within the outlier-enriched region on chromosome 10. In this region, we found reduced nucleotide diversity and a significant and negative Fu and Way’s H, which is sensitive to a high frequency of derived variants. These patterns--high differentiation, reduced diversity, as well as the same haplotype among individuals from resistant populations--indicates that a hard selective sweep of this region occurred across the four resistant populations. It also strongly suggests that this region contains at least some of the loci underlying glyphosate resistance in *I. purpurea*.

Intriguingly, we identified balancing selection among susceptible populations for this region on chromosome 10 (*i.e.* >2 Tajima’s D and Fu and Way’s H), which in this system would most likely be driven by crop rotations leading to herbicide on and off years, *i.e*., a pattern of alternating selection [43]. Further, and opposite our expectations, we found higher nucleotide diversity among resistant individuals for the outlier enriched regions found on chromosomes 1 and (to a lesser extent) 13. Such a pattern could be due to different loci responding to selection across resistant populations, or, and more likely, different haplotypes within resistant populations carrying the selected alleles. Unlike the dynamics we uncovered on chromosome 10, which suggest a hard selective sweep, the pattern of selection on chromosomes 1 and 13 are more aligned with a soft sweep model of evolution [43,44].

Within the five genomic regions enriched with outlier loci, we identified genes involved in the herbicide detoxification pathway, suggesting that glyphosate resistance is caused by herbicide metabolism in *I. purpurea*. The herbicide detoxification pathway is hypothesized to occur in three phases [11,45]: 1) activation, which is generally performed by cytochrome P450s, 2) conjugation, which is performed by GSTs or glycosyltransferases, and 3) transport into the vacuole, often by ABC transporters, which leads to the subsequent degradation of the herbicide. Multiple copies of each of these genes were present within the five outlier enriched regions. Within a 42.3 kb segment on chromosome 10, for example, we found seven duplicated, successive glycosyltransferase genes, with multiple non-synonymous SNPs present within the 1st, 4th, 5th and 7th glycosyltransferase genes. In addition to being present on the enriched region of chromosome 10, glycosyltransferases were also present within each of the other four outlier enriched regions. We likewise identified copies of ABC transporter and cytochrome P450 genes in two and three regions exhibiting selection, respectively. Although detoxification genes have yet to be functionally verified for glyphosate resistance in any weed species, transcriptomic surveys have shown that at least some of the genes involved in herbicide detoxification are associated with glyphosate resistance [28,46–48]. Additionally, we have previously shown that a cytochrome P450 transcript was up-regulated in artificially selected glyphosate resistant lineages of *I. purpurea* [28], supporting the conclusion that detoxification is a likely mechanism underlying glyphosate resistance in this species.

While our reduced representation population genomics and exome resequencing strategy has identified strong potential candidate genes associated with glyphosate resistance in *I. purpurea*, it is important to note that we found low levels of linkage disequilibrium between SNP markers (on average, r^2^ ∼0.03 across chromosomes). This suggests our initial reduced representation screen, which influenced the target exons we chose for exome resequencing, likely missed portions of the genome responding to selection from the herbicide. It also suggests, however, that the positive associations we did uncover (especially with our exome resequencing data) are likely to be loci that are involved in resistance, or very tightly linked to loci involved with resistance. Importantly, linkage was strongly elevated across outlier enriched regions compared to background levels of linkage for each of the chromosomes. These areas of increased linkage (defined as r^2^ > 0.25) in each outlier enriched region ranged between 84 kb to 3 MB in length, and support our findings of a genomic response to herbicide selection.

### Patterns of haplotype sharing across resistant populations suggests parallel and non-parallel responses to selection

Our initial population genomics screen across a genome-wide panel of ∼8K SNPs showed that resistant populations were not more related to one another than they were to susceptible populations, as would be expected under a scenario where resistance evolved in one lineage and moved *via* migration between locations. This, in addition to the ‘mosaic’ appearance of resistance among populations suggested that selection on standing variation was responsible for the repeated appearance of resistance in this species across the landscape. Another likely scenario, however, is one where migration introduced beneficial allele(s) that introgressed into the local background and then rapidly increased in frequency when exposed to very strong selection. The region under selection on chromosome 10 appears to follow this scenario. We found an identical haplotype within this region in high frequency across the resistant populations (>75% of individuals within populations with the same haplotype), and our formal test of convergence identified a very short standing time of the variant within this region (t = 5). Thus, the most likely model is one in which gene flow shared beneficial allele(s) between resistant populations which then started sweeping quickly and immediately, or within a few generations. This is likewise supported by our finding of low genome-wide patterns of linkage between SNPs, and evidence of a hard selective sweep, as indicated by low nucleotide diversity in this region and marks of positive selection indicating a high frequency of derived variants. Because this species employs a mixed mating system (*i.e.*, multilocus outcrossing rate (t_m_) = 0.5; [49]), it is plausible that resistance alleles, once introduced into the population, could quickly spread *via* outcrossing and then increase in frequency given strong selection.

Haplotypes from the other four outlier enriched regions were less consistently shared among the four highly resistant populations. The ‘resistant’ haplotype of the five outlier enriched regions were similar and in high frequency (>50%) in populations WG and SPC; genome-wide patterns of allele frequencies were also very similar between these two resistant populations (Fig 1C). This suggests that a highly similar resistant lineage is shared *via* migration between at least these two populations. The haplotypes of the other outlier enriched regions are in lower frequency among the other two resistant populations, BI and DW; further, pairwise F_ST_ values between resistant populations for the outlier enriched regions of chromosomes 1, 6, 13, and 15 were higher than the values among susceptible populations. These findings suggest a couple of possibilities: the presence of multiple haplotypes across these regions that carry resistance loci (*i.e.* soft sweep model of evolution), or the potential that resistance in this species is mostly attributable to the region on chromosome 10 that is shared and highly similar among resistant populations, with signs of selection from the other regions attributable to other factors. In support of the latter explanation, studies from other species have suggested that changes to a single step in the detoxification pathway are enough to provide some level of resistance [16]. However, coordinated upregulation of all of the genes from the detoxification pathway has been observed in grass species resistant to graminicide herbicides [50,51], suggesting that multiple components of this pathway are required for resistance. Unfortunately, there are few examples in which the genetic basis of NTSR resistance is known, making it difficult to draw conclusions on the importance of one gene versus the efforts of multiple genes in the detoxification pathway.

Interestingly, it is hypothesized that rather than detoxify the herbicides *per se*, these detoxification genes enable the plant to survive the resulting oxidative stress after being exposed to herbicide, a mechanism that may allow for resistance to several different herbicides [11]. This explanation--*i.e*. the ability to handle oxidative stress--could potentially underlie glyphosate resistance in *I. purpurea,* and further examination will be required to differentiate between the direct detoxification of glyphosate or an adaptive ability to respond to oxidative stress. Our results here, combined with that of previous work, also suggests the possibility of a slightly different story--a single gene (or set of them; *i.e.* region on chromosome 10) is enough to gain resistance but further involvement of other genes in the same pathway may lead to lower fitness costs. Individuals from the resistant BI population from TN, for example, share only the haplotype found on chromosome 10 in common with the other resistant populations. Interestingly, BI exhibits a higher cost of resistance than SPC and WG (26.9% germination vs 45.9% and 39.6%, respectively; [20]). This may indicate that loci specific to SPC and WG are important for ameliorating negative fitness costs of the changes in the chromosome 10 region.

## Conclusions

While there is strong evidence in support of genetic parallelism from cases of target-site resistance in other species [9,52], the genetic basis of non-target site resistance remains uncharacterized in most weeds [52,53]. Thus, we do not have a clear idea of the genetic mechanisms responsible for non-target site resistance, nor do we know how often the same mechanism is responsible for non-target site resistance across resistant lineages of the same weed. Our approach of using genome-wide scans and exome resequencing is an important step in understanding which broad-scale genetic changes may be responsible for resistance in *I. purpurea*, and whether or not the same genomic features respond to selection among populations.

Overall, our combined use of targeted sequencing, outlier analysis and exome re-sequencing provides a comprehensive view into the genomic basis of glyphosate resistance in a common and highly problematic agricultural weed. Our results suggest that genes responsible for herbicide detoxification are likely responsible for resistance in this species, with the important caveat that at this point we cannot determine if direct detoxification of the herbicide is occurring or if the species is able to respond to subsequent oxidative damage caused by the herbicide.Further, while we previously hypothesized that resistance across populations was due to selection on standing and shared genetic variation [4], the results we present here (stemming from the region on chromosome 10) support a scenario where gene flow between the resistant populations introduced the beneficial allele(s), followed by a hard selective sweep within a few generations. Finally, that we uncovered areas of genomic divergence among resistant populations within the regions showing signs of selection on chromosomes 1, 6, 13, and 15 suggests either different mutations/loci are involved with detoxification across populations, or that multiple haplotypes carrying the same adaptive alleles are responding to herbicide selection.

## Materials and Methods

### Seed collection and resistance phenotyping

Seeds were collected from populations across the range of *I. purpurea* (Table 1). In each population, seeds were sampled from multiple maternal individuals separated by at least 2 m. These seeds were used in a previously reported resistance assay to determine levels of resistance at field suggested rates of glyphosate [4].

### *EPSPS* sequencing

From the populations collected, we chose six high resistance (Avg. survival rate of populations, 84%) and five low resistance populations (Avg. survival rate of populations, 26%) that spanned the range of the collection in the U.S [4]. For each population we grew 2-5 (Avg. 4.1) plants from different maternal families in the greenhouse. Leaf tissue from each individual was collected and immediately frozen in liquid nitrogen. mRNA was extracted using the Qiagen RNeasy Plant kit and cDNA was created using Roche’s Transcriptor First Strand cDNA Synthesis Kit. Primers were designed based on *Convolvulus arvensis EPSPS* (GenBank: EU698030.1). These primers were used in a PCR to amplify the *EPSPS* coding regions using Qiagen’s Taq PCR Master Mix kit, followed by cleaning using GE’s Superfine Sephadex. Samples were then Sanger sequenced by the sequencing core at the University of Michigan. Bases were scored usingPHRED [54] followed by visual confirmation of heterozygous sites. Each of the copies of the *EPSPS* were aligned across all individuals using MafftWS [55] *via* Jalviewer [56] (Genbank: MK421977-MK422097). Variable sites were identified and used to obtain allele frequencies for the pool of resistant and susceptible populations separately. We used a χ^2^ test to determine if allele frequencies varied between resistant and susceptible populations, and likewise determined if allele frequencies were correlated with population-level resistance values using Pearson’s correlation. P-values were adjusted for multiple tests using the Benjamini and Hochberg [57] correction. We also calculated observed and expected heterozygosity using adegenet [58,59] and tested for Hardy-Weinberg equilibrium using 1000 bootstraps in pegas [60]. To compare to other known *EPSPS*, we downloaded several protein sequences from GenBank and aligned them to our translated amino acid sequences using tCoffee [61] in Jalview [56] (S1 Fig).

### SNP genotyping

Eight populations were chosen to investigate non-target site resistance: 4 low resistance (Avg. survival rate, 16%) and 4 high resistance populations (Avg. survival rate, 89%) (Fig 1A; Table 1, data from [4]). Seeds from up to 10 maternal families per population were germinated and leaves were collected and frozen for DNA extractions. A total of 80 individuals were used for SNP genotyping.

DNA was extracted using a Qiagen Plant DNeasy kit. Genomic DNA was converted to nextRAD sequencing libraries (SNPsaurus). The nextRAD method for GBS (genotyping-by-sequencing) uses a selective PCR primer to amplify genomic loci consistently between samples; nextRAD sequences the DNA downstream of a short selective priming site. Genomic DNA (7 ng) from each sample was first fragmented using a partial Nextera reaction (Illumina, Inc), which also ligates short adapter sequences to the ends of the fragments. Fragmented DNA was then amplified using PhusionÂ® Hot Start Flex DNA Polymerase (NEB), with one of the Nextera primers modified to extend eight nucleotides into the genomic DNA with the selective sequence TGCAGGAG. Thus, only fragments starting with a sequence that can be hybridized by the selective sequence of the primer were amplified by PCR. The 80 dual-indexed PCR-amplified samples were pooled and the resulting libraries were purified using Agencourt AMPure XP beads at 0.7x. The purified library was then size selected to 350-800 base pairs. Sequencing was performed using two runs of an Illumina NextSeq500 (Genomics Core Facility, University of Oregon). This resulted in 42,004,808,475 bp total, with an average of 525,060,106 bp per individual (Genbank: XXXX).

To control for repetitive genomic material or off-target or error reads, coverage per locus was determined using reads from 16 individuals and loci with overly high or low read counts were removed (*i.e.* above 20,000 or below 100). The remaining reads were aligned to each other using BBMap [62] with minid = 0.93 to identify alleles, with a single read instance chosen to represent the locus in a pseudo-reference. This resulted in 263,658 loci. All reads from each sample were then aligned to the pseudo-reference with BBMap and converted to a vcf genotype table using Samtools.mpileup (filtering for nucleotides with a quality of 10 or better), and bcftools call [63]. The resulting vcf file was filtered using vcftools [64]. SNPs were removed if there was a minimum allele frequency less than 0.02, a read depth of 5 or less, an average of less than 20 high quality base calls or more than 20% of individuals exhibited missing data. This left 8,210 SNPs.

### RAD-seq analysis

Basic population genetic statistics (He, Ho, and F_IS_) were calculated *via* poppr [65] and hierfstat [66] packages and can be found in S2 Table. fastStructure [67] was used to detect population structure (S2 Fig). A PCA analysis on individual allele frequencies was used to investigate structure using the dudi.pca function in the adegenet package [58,59] in R. Tassel was used to construct a neighbor-joining tree from pairwise relatedness [68]. Bootstraps of loci were conducted using a custom script, with 500 replications.

We used two outlier-based programs to identify potential loci under selection. We first used BayeScan (version 2.1, [69], which assumes an ancestral population from which each sampled population differs by a given genetic distance. Pairwise F_ST_ values are calculated between each sampled population and the ancestral population, thus correcting for differences in population structure. These F_ST_ values are then used in a logistic regression that includes a population specific factor (the structure across all loci) and a locus specific factor. If the locus-specific factor significantly improved the model, it implies that something abnormal is occurring, which is assumed to be natural selection. We used the default settings (false discovery rate of 0.05) to identify loci that showed evidence of high F_ST_ between the resistant and susceptible populations.

The second program, bayenv2, identifies correlations between locus specific allele frequencies and an environmental variable [70,71]; in our work, the “environment” is the level of resistance per population. This program uses “neutral” loci to create a genetic correlation matrix against which each SNP is tested for a correlation between its frequency and the environment. In essence, the allele frequencies are modeled based on solely the neutral correlation matrix and with the addition of the environmental variable. Loci potentially under selection are then identified using the Bayes Factor (the support for the model with the environmental variable added) and the Spearman’s correlation coefficient. To estimate the “neutral” population structure, we removed any SNPs from sequences that aligned (*via* bowtie) with either the *I. purpurea* or *Lycium sp.* (from 1kp data, [72] transcriptome (only 35% of SNPs aligned to either) and then used the final matrix outputted from the correlation matrix estimation after 100,000 iterations. All SNPs were then run with the environment being either −1 for the susceptible populations or 1 for the resistant populations, and a burn-in of 500,000 with a total of 5 runs was performed (correlation between runs was >0.80). Following Gunter & Coop [71], we identified outlier loci with the highest 1% of Bayes Factors and the 5% most extreme Spearman correlation coefficients averaged over the 5 runs.

We compared pairwise F_ST_s for the resistant and susceptible populations using the full data set and the outlier data set using 4P [73]. We calculated Weir and Cockerham [32] pairwise F_ST_ values for each data set (overall SNPs and outlier SNPs) for each pair of populations to calculate the average F_ST_ among resistant populations and susceptible populations. To obtain 95% confidence intervals around these estimates, we performed the same steps on 1000 randomly selected sets of loci (sampled with replacement).

### *De novo* genome assembly for exome resequencing

We annotated RAD-seq outliers by using a BLASTN analysis to align them to a draft genome sequence from highly homozygous *I. purpurea* individual. To generate the draft genome sequence, DNA from a single individual was sequenced using PacBio (11 SMRT cells) and Illumina (2 lanes of 100 bp, paired end) sequencing (Genbank: XXXX). PacBio reads were filtered for adaptors and to remove low quality (<0.8) and short read lengths (<500 bp). Illumina reads were trimmed of low quality sequences using trimmomatic [74]. Illumina reads were assembled using ABYSS-PE k=64 [75]. This resulted in 1,933,851 contigs with lengths ranging from 64-94,907 bp (N50=6,790) for a total of 631,125,096 bp. LoRDEC [76] was used to error correct the PacBio sequences using the raw Illumina reads followed by trimming of weak regions. This resulted in 4,621,037 reads and 1,823,002,799 bp. These sequences were then combined with the Illumina assembled contigs using DBG2OLC (k=17, kmer coverage threshold = 2, min overlap = 10, adaptive threshold = 0.001, LD1=0) [77]. This resulted in 17,897 scaffolds with lengths ranging from 231-162047 bp (N50=15,425) for a total of 194,708,849 bp. This PacBio+Illumina assembly as well as the Illumina-alone assembly were used in a BLASTN analysis with each of the RAD-seq outliers. For those with genomic hits, putative genes on the contig were determined using AUGUSTUS [78], FGenesH [79], SNAP [80] and tRNAScan [81], which were used to design target capture probes.

### Target capture exome re-sequencing

We next designed probes for exome sequencing of loci that were either identified from our population genomics RAD-seq screen or loci have been shown to correlate with resistance in other species. We used a variety of methods to select possible capture probe sequences. First, we used a BLASTN [82] analysis to select transcripts matching our RAD-seq outliers -we BLASTed the 75 bp of the RAD-seq tags that contained outlier SNPs from either the BayeScan or bayenv2 analyses using the full dataset against transcripts in an *I. purpurea* transcriptome [28] and selected the top hit for each (30 transcripts, min e-value = 3e-7). Second, we selected transcripts that were previously identified as differentially expressed in an RNAseq experiment[28] which compared artificially selected resistant and susceptible lines following herbicide application (19 sequences). Third, we used the two EPSPS mRNA sequences (2 sequences). Fourth, we used a BLASTN analysis to select the putative genes on genomic contigs that matched our outlier SNPs. We BLASTed 75 bp of the RAD-seq tags that contained outlier SNPs to the two draft *I. purpurea* genomes described above and then selected the resulting coding sequences from the putative genes (171 sequences, min e-value=7e-14). Additionally, we randomly chose an even number of transcripts from the transcriptome to serve as our controls (214 sequences).

These 436 sequences were then used to design the capture probe candidates. Candidate bait sequences were 120 nt long, with a 4x tilling density. Each bait candidate was BLASTed against the *I. trifida* genome [83], and a hybridization melting temperature (Tm)* was estimated for each hit. Non-specific baits were filtered out (Additional candidates pass if they have at most 10 hits 62.5 – 65°C and 2 hits above 65°C, and fewer than 2 passing baits on each flank.) This led to 16,078 baits, with a total length of 580,421 nt.

To generate material for sequencing, five seeds from each of the 8 populations used in the previous population genomics screen (Fig 1A, Table 1) were grown in the greenhouse, leaves were collected, and DNA was extracted from young leaf tissue using a Qiagen Dneasy Plant Mini kit. Genomic DNA was sent to MYcroarray for library preparation and target enrichment using the MYbaits (R) system. Genomic DNA was sonicated and bead-size-selected to roughly 300nt fragments, which was then used to create libraries using the Illumina (R) Truseq kit. A total of 6 cycles of library amplification using dual-indexing primers was applied, and index combinations were chosen to avoid potential sample misidentification due to jumping PCR during pooled post-capture amplification [84]. Pools of 3 or 4 libraries each were made, combining 200 or 150 nanograms of each library, respectively. These pools were then enriched with our custom MYbaits (R) panel (following the version 3.0 manual). After capture cleanup, the bead-bound library was amplified for 12 cycles using recommended parameters, and then purified with SPRI beads. These amplified, enriched library pools were combined in proportions approximating equimolar representation of each original library and sequenced on 2 lanes of Illumina 4k 150 PE. Our coverage goal was >30x depth per individual per locus.The resulting sequences were trimmed of adaptor sequences and low quality bases using cutadapt (q<10 removed). On average we sequenced 11.6 million reads (min = 5.9 million, max = 13.8 million) for each individual (Genbank: XXXX).

We next assembled the sequenced reads into contigs to perform SNP calls. We used trimmed sequences from one individual and default settings in Megahit [85] to assemble reference contigs (24,524,768 reads assembled into 67,266 contigs; N50 = 458 bp; range 200-16167 bp; S3 Table). For each individual, trimmed sequences were aligned to the assembled contigs using bwa [86], and SNPs were called and then filtered using the GATK pipeline ([87– 89]; overview of process: variants were initially called, individuals jointly genotyped, bases recalibrated based on filtered initial variants, and variants were recalled and jointly genotyped; specific commands: QD<2.0, FS>60, MQ<40, MQRankSum <12.5, RedPosRankSum<-8, minimum allele frequency >0.02, min mean depth > 5, max missing <0.8, min Q >20). After examining coverage per site, we found several contigs to have extremely high coverage and nearly 100% heterozygosity, suggesting multiple sequences were collapsed into 1 variant. Thus, we eliminated sites that had greater than 80% heterozygosity across individuals, or had more than twice the average coverage. This left 152,636 SNPs on 26,988 contigs for downstream analyses (N50 = 530; range 200-16167 bp). Bwa [86] was used to align the *de novo* contigs to the probes to estimate the percentage of SNPs that were from target capture probes. Fifty-one percent of these SNPs exhibited significant homology to one of the original probe sequences. In addition to analyzing these contigs, we also examined the contigs that did not exhibit significant homology to the probe sequences (hereafter non-probe contigs). The coverage for non-probe contigs was lower than that for probe contigs (23x average vs 33x), however because 13x coverage is sufficient to call heterozygous SNPs in a diploid [90] we chose to use both to increase our sampling of the genome. To annotate the contigs, we used a local TBLASTX analysis against Arabidopsis cDNA (from TAIR: [91]; e-value 0.001, and chose the sequence with the highest e-value for identification).

### Outlier analysis of targeted exome re-sequencing

We used BayeScan and bayenv2 as above to identify putative adaptive loci from the targeted exome re-sequencing. To reduce the effect of linkage among loci, we randomly chose 1 SNP per 1000 bp using vcftools (27,225 SNPs retained). To estimate the neutral population structure for bayenv2, 2000 contigs were randomly selected from contigs that did not map to the probes designed for the outliers.

To place the SNPs into a genomic context we aligned them to the *I. nil [29]* genome using BLAT [92] and liftover [93]. A total of 124,149 SNPs aligned to the genome. By visual analysis we identified five regions with a large majority of significant outliers (*i.e.* 71% of outlying SNPs). We delimited each of these ‘enriched regions’ by the first and last outlier of each region, and searched the regions for genes involved in non-target site resistance using the following GO terms and gene names: GO:0009635, GO:0006979, GO:0055114, glycosyltransferase, glutathione s-transferase, ABC transporters and cytochrome P450s. We randomly selected 5 regions of the same size from the *I. nil* genome and counted the number of genes from the above gene families to determine if outlier enriched regions contained more of these genes of interest than expected due to chance. We repeated this 1000 times to create an empirical distribution, which was then used to determine the percentile of the observed data. We next determined if outliers outside of the enriched regions were more, less, or equally likely to be located near a gene family of interest (*i.e*., glycosyltransferase, ABC transporter, etc). To do so, we counted the number of genes of each family within ∼4MB (the largest of the 5 regions) from outliers found outside the five enriched regions and then compared the distributions of these outliers to those found within the enriched regions. We used CooVar [94] with the *I. nil* gene models to predict the protein level changes for each SNP that aligned to the *I. nil* genome and to determine if SNPs were from nonsynonymous or synonymous sites.

For estimates of genetic differentiation and diversity, we calculated G_ST_ [95], nucleotide diversity (as the ratio of susceptible to resistant individuals; piS/piR), Tajima’s D [30], and Fu and Way’s H [31] over 25 SNPs windows using customized scripts from [96]; 124,149 SNPs in analyses). Additionally, we used vcftools to calculate pairwise F_ST_ estimates [32] among the resistant and susceptible populations separately for each SNP. Negative F_ST_ values will occur when there is little genetic variation, and thus we set any negative value to zero. We then compared the average pairwise F_ST_ of the resistant populations versus the susceptible populations within the 5 outlier enriched regions.

We used hierarchical modeling to determine if the resistant populations had similar haplotype structure in outlier containing regions of the genome, which would potentially indicate that resistance is controlled by the same genetic basis across populations. We grouped sequenced individuals into either those that exhibited the putative susceptible allele (‘S’ group) or the putative resistant allele (‘R group’) for each contig. To do so, the pairwise genetic distance between each individual was calculated based on all SNPs in each contig using the dist.gene command from the ape package (vers 5.0; [97]) in R [98]. This genetic distance matrix was reduced to 2 dimensions by multidimensional scaling using the cmdscale and eclust commands [99]. These two dimensions were then used to hierarchically cluster the populations into 2 groups using kmeans clustering. The group that contained less than half of the individuals from the susceptible populations was deemed the ‘R’ group (*i.e.* those that are genetically different from the majority of the susceptible individuals and presumably have the allele that aids in resistance).

### Linkage

We used the exome resequencing data to examine patterns of linkage across the genome and within the outlier-enriched regions. We estimated LD as the correlation coefficient (r^2^) between each SNP pair using the program GUS-LD (genotyping uncertainty with sequencing data-linkage disequilibrium; [100]), a likelihood method developed to estimate pairwise LD using low-coverage sequencing data. GUS-LD controls for under-called heterozygous genotypes and sequencing errors, which are a known problem with reduced representation sequencing. We estimated LD for each chromosome that exhibited an outlier enriched region (chr1, chr6, chr10, chr13, chr15) using the SNPs identified across all individuals using the exome resequencing dataset. We used only biallelic SNPs of at least 2% frequency and with <20% missing genotype calls since rare alleles can influence the variance of LD estimates. Only SNPs that could be aligned to the *I. nil* genome were used in the analysis, and we used the reduced SNP dataset (1 SNP/kb) to reduce processing time (∼10K SNPs used overall). The number of SNPs used per chromosome ranged from 1189 to 3191 and are presented in S5 Table. Linkage decay was not estimated due to the granular nature of the data; instead, we report r^2^ values averaged over the entire chromosome as a background estimate of LD, along with the 3rd quartile of r^2^ values, and the average of r^2^ values of SNPs located within 10 kb of one another.

### Test of convergence

Given evidence that the outlier-containing region on chromosome 10 showed the strongest sign of differentiation between the resistant and susceptible populations (see Results), we applied the inference method of Lee and Coop (2017) to examine the most likely mode of adaptation within this region. This composite likelihood based approach, explained in full in [101] both identifies loci involved in convergence and distinguishes between alternate modes of adaptation--whether adaptation is due to multiple independent origins, if adaptive loci were spread among populations *via* gene flow, or were shared among populations due to selection on standing ancestral variation. We first estimated an F matrix to account for population structure using SNPs from scaffolds on Chr 3, 7, and 14 that showed no evidence of selection from our outlier analyses (S6 Table). We then used all SNPs (N = 2248) on a scaffold from the *I. purpurea* assembly (that aligned to *I. nil* scaffold BDFN01001043) to apply the inference framework to the region on chromosome 10 that exhibited signs of selection. We estimated the maximum composite likelihood over a grid of parameters used to specify these models (S7 Table). We allowed two of the resistant populations (WG and SPC) to be sources of the variant in the migration model. Additionally, and following [101], we used an N_e_ = 7.5 × 10^5^.

### Functional annotation of nontarget site resistance genes

To predict the putative function of genes within the five enriched regions, we used a BLASTN analysis to generate a network graph of each of our target gene families (cytochrome P450s [102], glycosyltransferases [103], and ABC transporters [104]) based on homology to *Arabidopsis thaliana* genes. We used the *I. nil* genes from the outlier regions and all *A. thaliana* genes from each gene family in an all-by-all BLASTN, with an e-value = 1^−10^ for the cytochrome P450s and ABC transporters. For the glycosyltransferases, we first used a conserved domain search to identify glycosyltransferase genes in the 5 outlier-enriched regions and used these in the BLASTN search, with an e-value cutoff of 1^−1^ due to very low conservation among genes within this family, a widely-recognized problem [105]. The resulting bit-score of the BLASTN analysis were then used in cytoscape [106] to visualize the relationships, with colors denoting the families (P450s and ABC transporters) or conserved coding domains (glycosyltransferases).

### Accession numbers

*EPSPS* sequencing data (MK421977-MK422097), NextRAD sequencing data (XXXX), genome assembly (XXXX) and Exome resequencing data (XXXX) are available in GenBank.

## Supporting Information

**S1 Fig.**
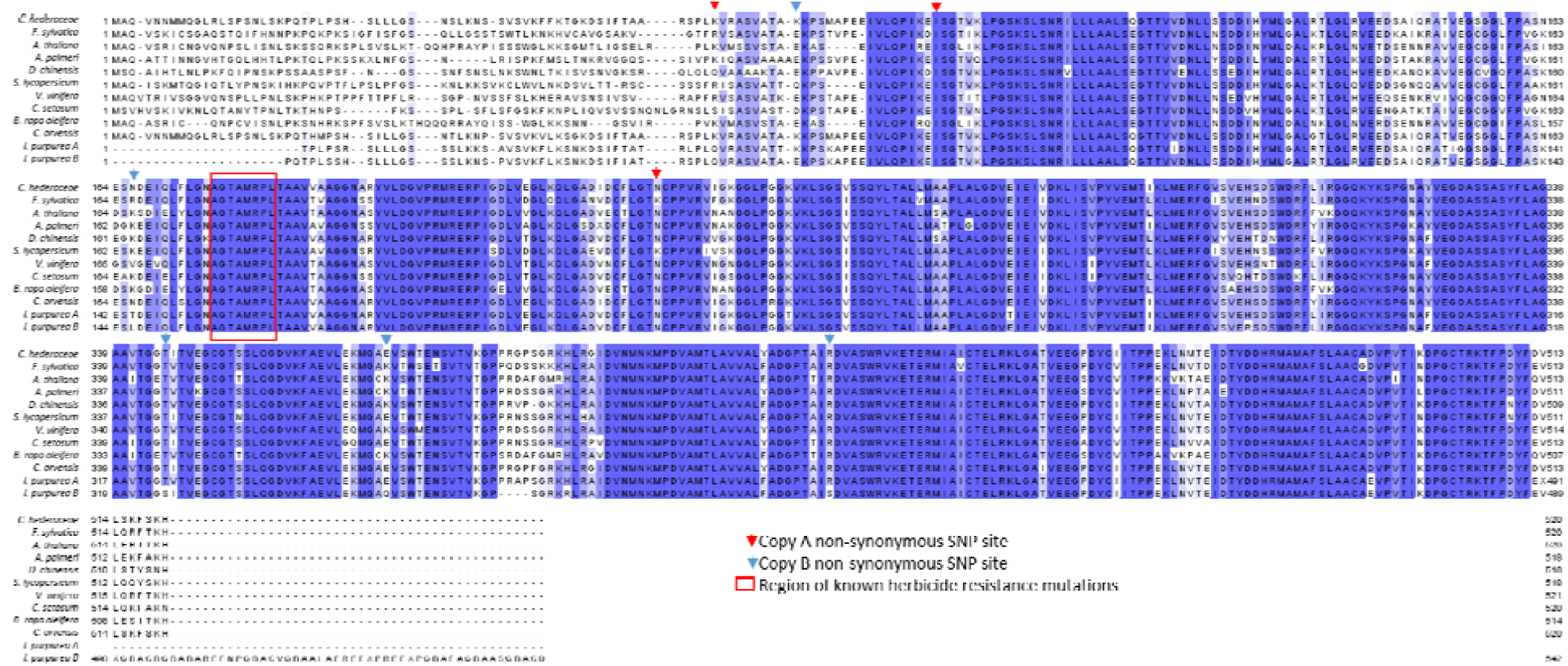
No sequence differences in *EPSPS*. Comparison of amino acid sequences of *Ipomoea purpurea* EPSPS protein sequence (bottom two sequences) with other reported EPSPS proteins in the NCBI database (gi|170783792, gi|76782198, gi|15225450, gi|257228989, gi|16751569, gi|460388790, gi|225454012, gi|374923051, gi|46095337, gi|189170087) shows no sequence variation within the region known to affect herbicide resistance (inside red outline). Non-synonymous changes (red and blue arrows) outside of this region likewise do not correlate with resistance (Table S1).

**S2 Fig.**
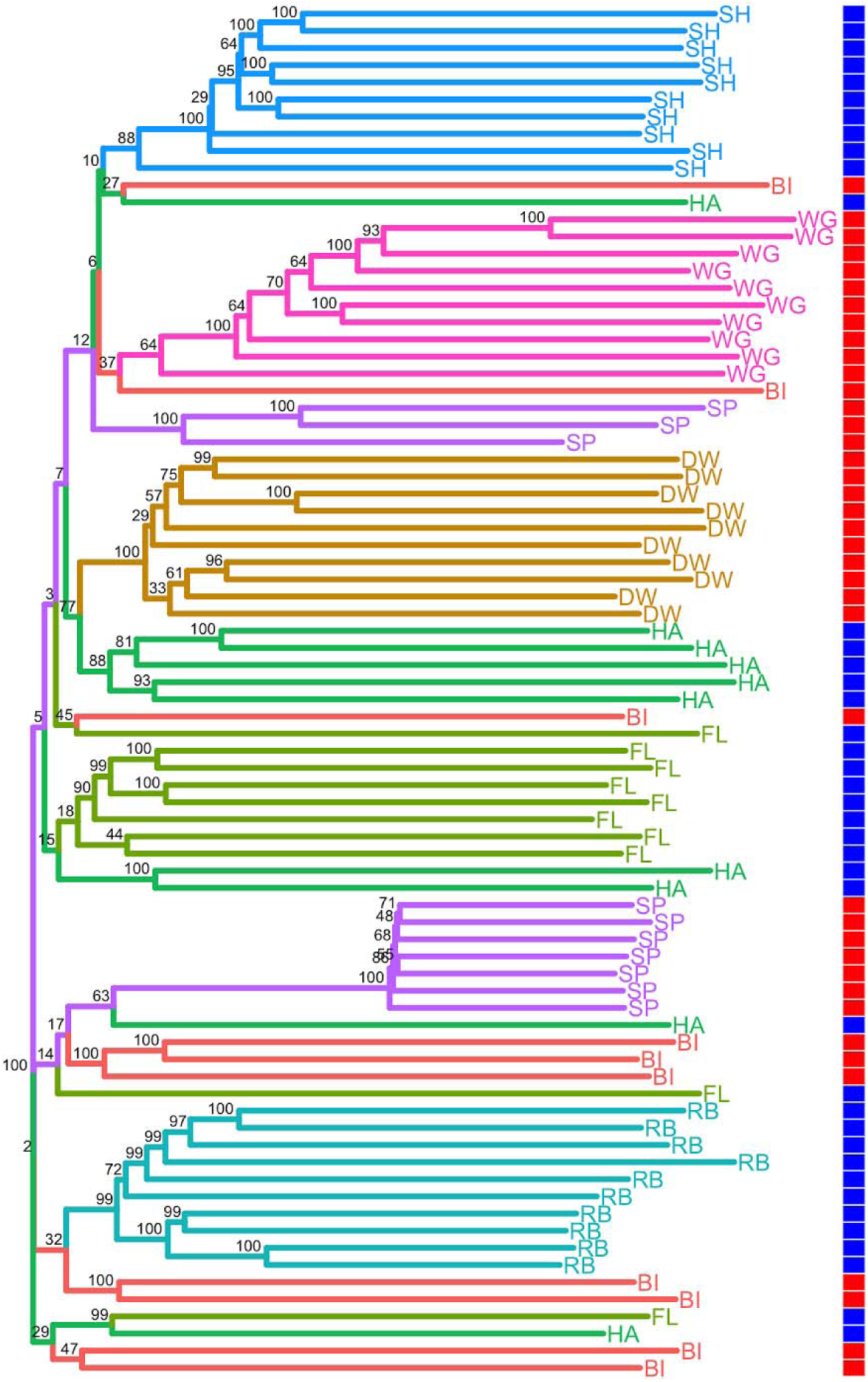
Neighbor joining tree using all of the RAD-seq SNP loci. On the tree, populations are denoted by color and tip labels; values at nodes are percent bootstrap support; population level resistance is denoted by the color in the column (red=resistant, blue=susceptible).

**S3 Fig.**
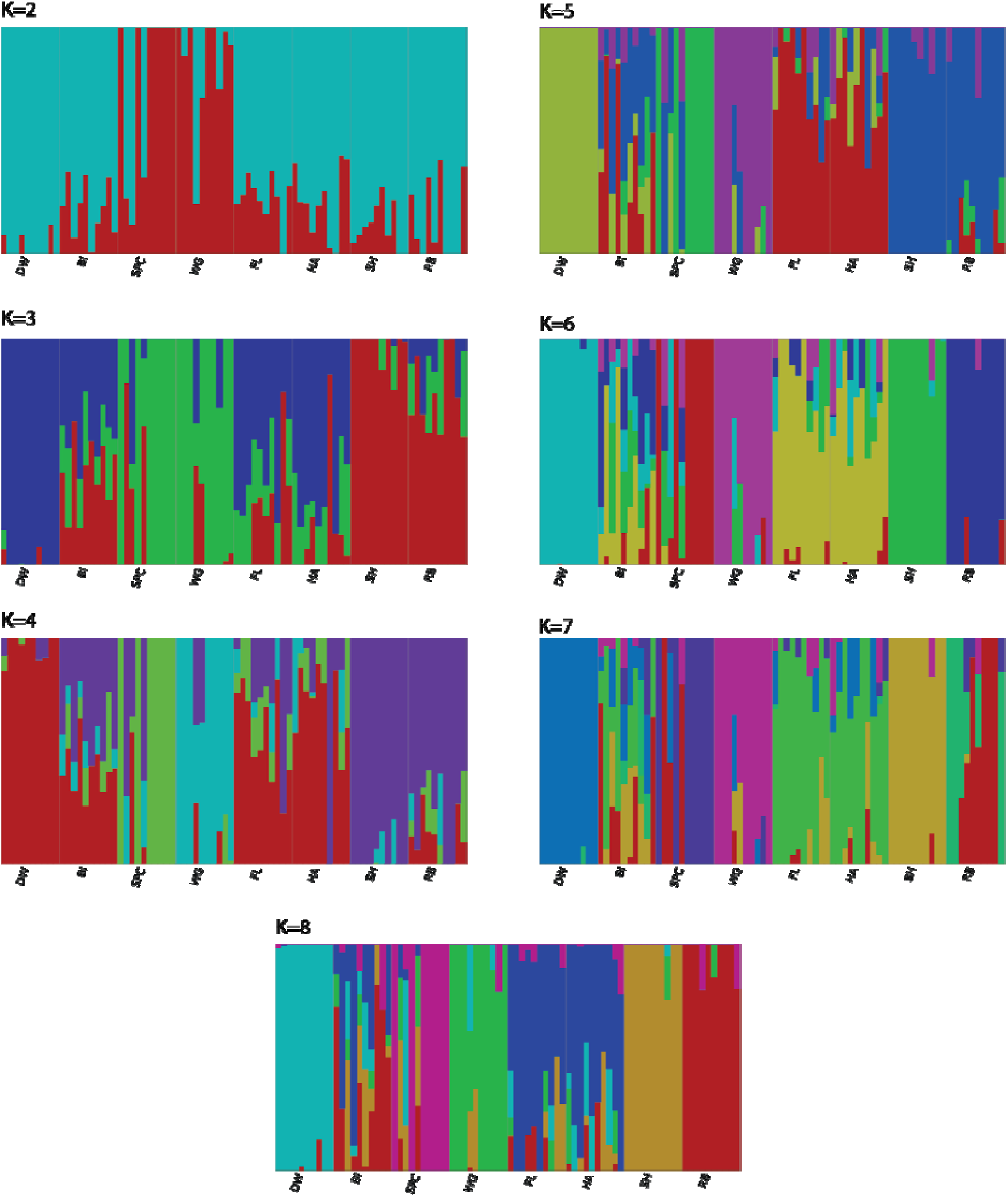
Population structure analyses. At K=2 FastStructure results for the RADseq data do not show the resistant populations (first four populations on the left) segregating into a distinct group, suggesting they are not from a single origin. FastStructure analysis suggests either K=6 or K=7 as the best model, both of which leads to some populations being highly admixed (*e.g.* BI) while others are fairly homogenous (*e.g.* SH).

**S4 Fig.**
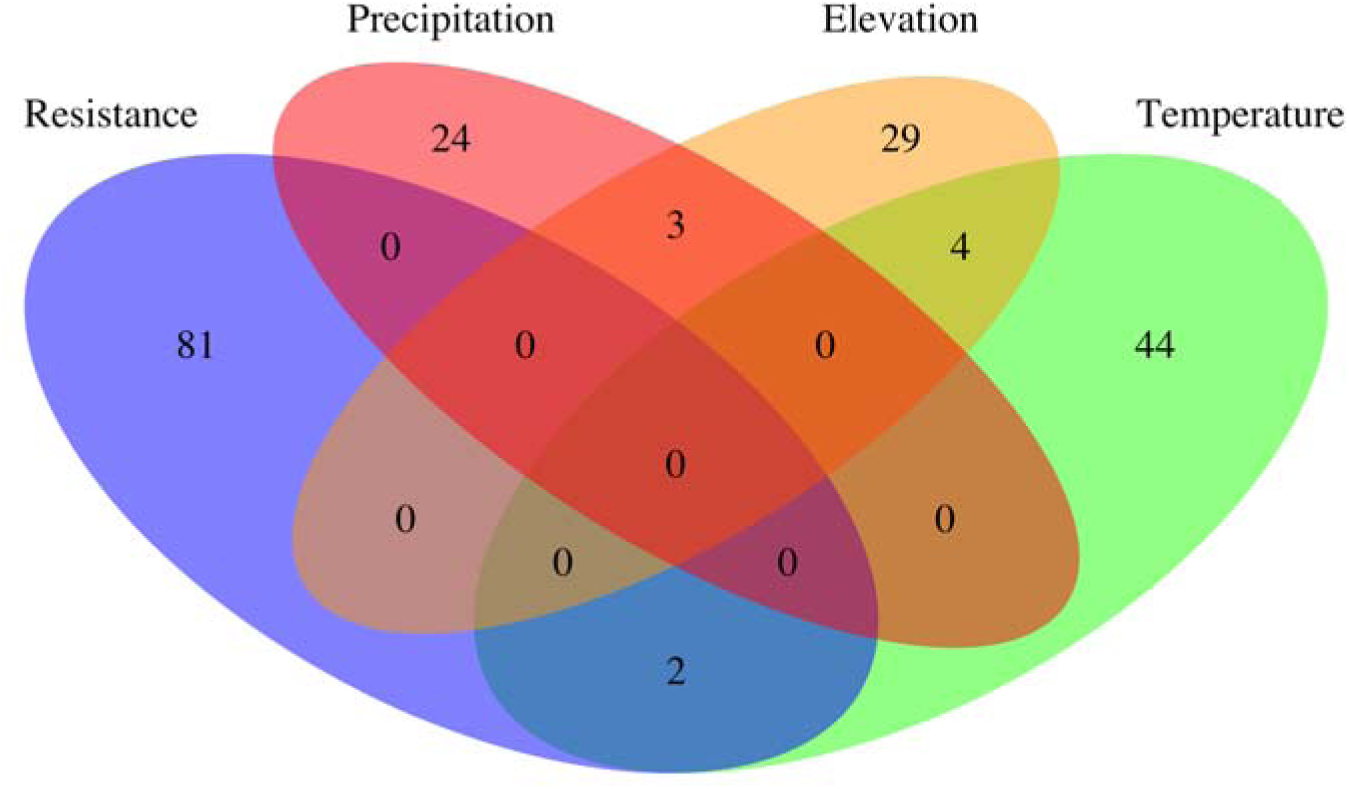
RADseq outliers associated with environmental variables. Based on BayEnv2 analyses using environmental variables, we identified 50 loci that correlated with minimum temperature of coldest month, only 2 of which overlapped with the resistance outliers; 27 loci correlated with precipitation of the driest month, 0 of which overlapped with the resistance outliers; 36 loci correlated with elevation, 0 of which overlapped with the resistance outliers.

**S5 Fig.**
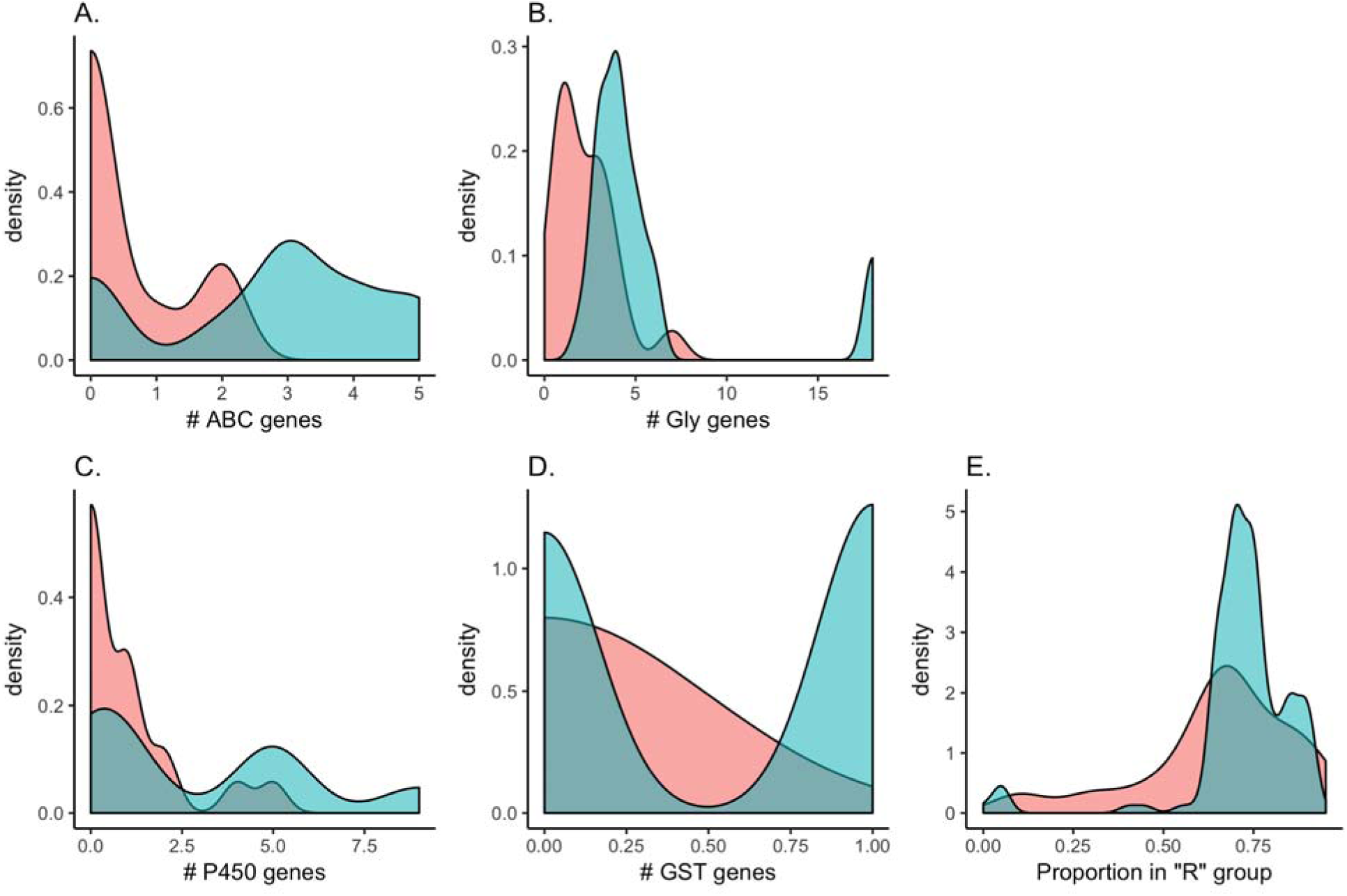
Differences between outliers inside and outside of outlier enriched regions. (A-D) Distributions of the number of genes within 4 mb of an outlier, either inside (blue) or outside (red) an outlier-enriched region. For each type of gene, the outliers outside of the regions show a left-skewed distribution indicating fewer close detoxification genes for (A) ABC transporters, (B) Glycosyltransferases, (C) Cytochrome P450s and (D) Glutathione S-transferases. (E) Outliers outside of the regions have lower frequencies of the resistant haplotype than those inside the regions, suggesting they are more population specific.

**S6 Fig.**
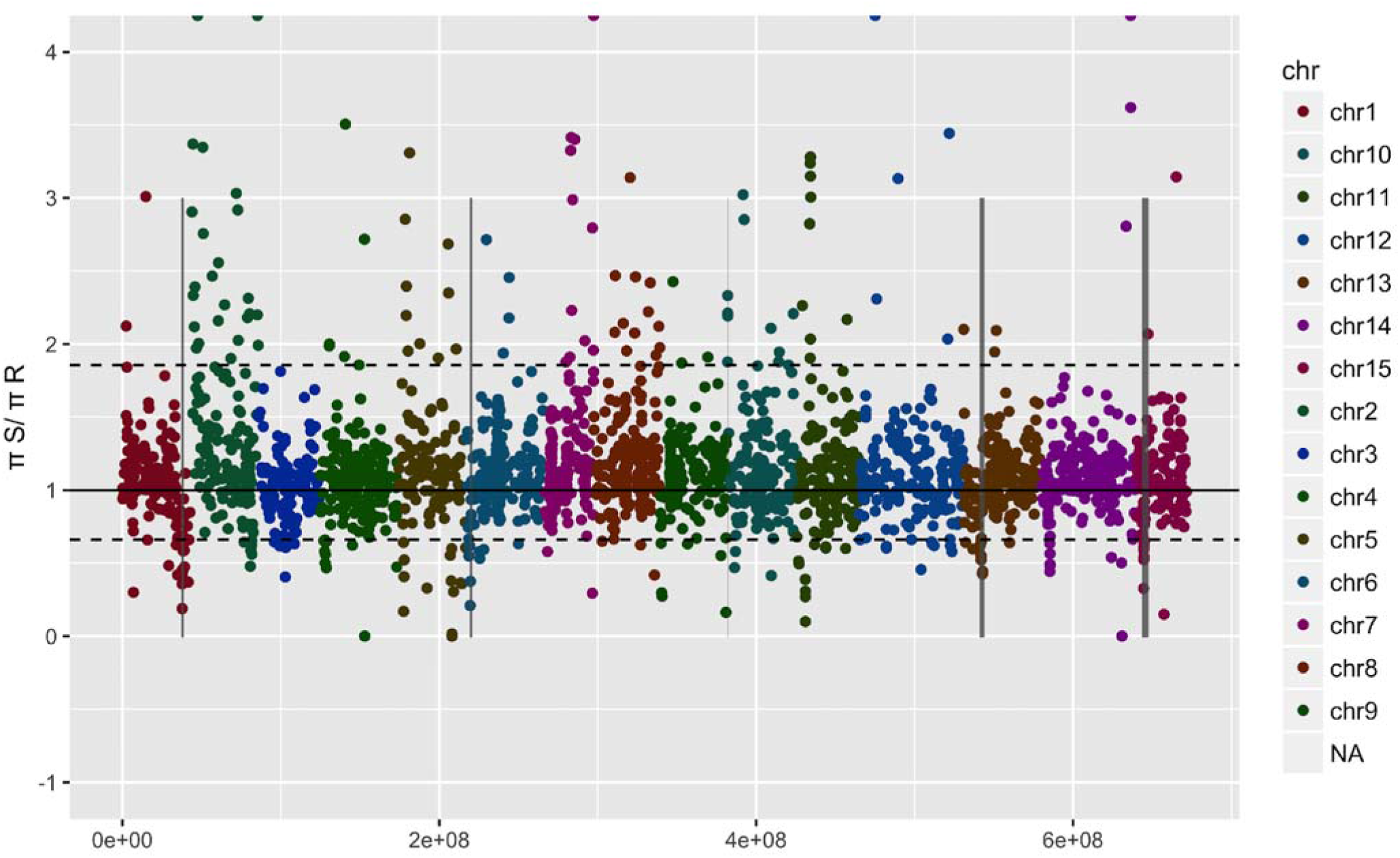
Nucleotide diversity across all SNPs that aligned to the *I. nil* genome. Data are shown are the ratio of susceptible to resistant individual nucleotide diversity. Grey bars indicate the outlier enriched regions identified on chromosomes 1, 6, 10, 13, and 15. Dashed lines show the 5% most extreme genome-wide values.

**S1 Table.** *EPSPS* SNP data for gene copy A and B. SNP #=the location of the SNP after alignment with *EPSPS* from *Convulvulus arvensis*, Ho=observed heterozygosity (across all samples), He=expected heterozygosity, HWE p-value=p-value for test of Hardy-Weinberg equilibrium from permutation test, Alleles=SNP alleles, Syn=whether a synonymous change (as determined by alignment with *C. arvensis* sequence), P-value chi-squared R vs S=p-value for test of allele frequency difference between resistant and susceptible populations, P-value cor with resistance=adjusted p-value for correlation with survival.

| SNP # | Ho | He | HWE p-value | Alleles | Syn | P-value chi-squared R vs S | P-value cor with resistance |
| --- | --- | --- | --- | --- | --- | --- | --- |
| <u>EPSPS A</u> |  |  |  |  |  |  |  |
| 102 | 0.19 | 0.47 | 0 | A:G | Y | 0.68 | 0.20 |
| 188 | 0.02 | 0.02 | 1 | A:G | N | 0.89 | 0.45 |
| 234 | 0.2 | 0.47 | 0 | C:G | Y | 0.68 | 0.19 |
| 247 | 0.2 | 0.47 | 0 | C:T | Y | 0.68 | 0.19 |
| 265 | 0.02 | 0.02 | 1 | A:G | N | 0.89 | 0.45 |
| 689 | 0.11 | 0.11 | 1 | A:T | N | 0.77 | 0.12 |
| 690 | 0.11 | 0.11 | 1 | C:A | N | 0.77 | 0.12 |
| 741 | 0.16 | 0.48 | 0 | G:A | Y | 0.64 | 0.19 |
| 831 | 0.18 | 0.47 | 0 | C:T | Y | 0.66 | 0.19 |
| 936 | 0.2 | 0.48 | 0 | T:C | Y | 0.66 | 0.20 |
| 1194 | 0.21 | 0.48 | 0.001 | T:C | Y | 0.69 | 0.21 |
| 1425 | 0.18 | 0.49 | 0 | A:G | Y | 0.67 | 0.19 |
| 1500 | 0.11 | 0.5 | 0 | T:G | Y | 0.57 | 0.19 |
| 1503 | 0.14 | 0.5 | 0 | T:C | Y | 0.62 | 0.19 |
| <u>EPSPS B</u> |  |  |  |  |  |  |  |
| 214 | 0.18 | 0.42 | 0.001 | G:A | N | 0.85 | 0.59 |
| 246 | 0.18 | 0.4 | 0 | T:G | Y | 0.84 | 0.59 |
| 372 | 0.18 | 0.42 | 0.001 | C:T | Y | 0.80 | 0.59 |
| 496 | 0.18 | 0.42 | 0 | C:A | N | 0.85 | 0.59 |
| 497 | 0.27 | 0.63 | 0 | T:C:G | N | 0.89 | 0.17 |
| 606 | 0.18 | 0.4 | 0.001 | T:G | Y | 0.86 | 0.59 |
| 666 | 0.18 | 0.4 | 0 | T:C | Y | 0.84 | 0.59 |
| 729 | 0.23 | 0.5 | 0.001 | T:A | Y | 0.89 | 0.45 |
| 792 | 0.16 | 0.31 | 0.002 | C:T | Y | 0.69 | 0.17 |
| 921 | 0.16 | 0.34 | 0.003 | C:T | Y | 0.67 | 0.45 |
| 963 | 0.16 | 0.34 | 0.005 | C:T | Y | 0.67 | 0.45 |
| 1029 | 0.3 | 0.49 | 0.01 | A:G | Y | 1.00 | 0.45 |
| 1034 | 0.3 | 0.49 | 0.007 | G:C | N | 1.00 | 0.45 |
| 1044 | 0.3 | 0.49 | 0.012 | T:C | Y | 1.00 | 0.45 |
| 1053 | 0.18 | 0.35 | 0.002 | C:T | Y | 0.67 | 0.17 |
| 1114 | 0.3 | 0.49 | 0.012 | C:G | N | 0.98 | 0.45 |
| 1278 | 0.02 | 0.49 | 0 | T:A | N | 0.97 | 0.48 |
| 1359 | 0.16 | 0.34 | 0.002 | C:G | Y | 0.67 | 0.45 |
| 1392 | 0.3 | 0.49 | 0.011 | A:G | Y | 1.00 | 0.45 |
| 1401 | 0.3 | 0.49 | 0.022 | G:A | Y | 1.00 | 0.45 |
| 1413 | 0.16 | 0.34 | 0.001 | A:G | Y | 0.67 | 0.45 |
| 1503 | 0.02 | 0.02 | 1 | C:T | Y | 0.89 | 0.66 |

**S2 Table.** Population genetics parameters for the RADseq SNPs. Population = population abbreviation. Ave N/locus = average number of individuals with high quality allele data per locus. % loci missing = average percent of the population with missing data per locus. Ho = observed heterozygosity. He = expected heterozygosity. FIS = Wright’s inbreeding coefficient.

| Population | Ave N/locus | % loci missing | Ho | He | FIS |
| --- | --- | --- | --- | --- | --- |
| RB | 9.518758 | 0.048124 | 0.239929 | 0.260012 | 0.101951 |
| HA | 9.585627 | 0.041437 | 0.276958 | 0.325516 | 0.135513 |
| BI | 9.223264 | 0.077674 | 0.282875 | 0.335554 | 0.151809 |
| DW | 9.539342 | 0.046066 | 0.242219 | 0.267021 | 0.112749 |
| FL | 9.610597 | 0.03894 | 0.278797 | 0.309048 | 0.107022 |
| SH | 9.648599 | 0.03514 | 0.26598 | 0.247444 | -0.03845 |
| SPC | 9.632156 | 0.036784 | 0.207493 | 0.222291 | 0.119612 |
| WG | 9.488307 | 0.051169 | 0.209562 | 0.251296 | 0.209932 |

**S3 Table.** Assembly statistics for the Illumina genome assembly (using ABYSS-PE), the PacBio + Illumina genome assembly (using DBLOG2), the resequencing assembly (using Megahit) and the resequencing assembly contigs containing SNPs.

|  | Illumina | PacBio+Illumina | Denovo contigs | Denovo contigs with SNPs |
| --- | --- | --- | --- | --- |
| Number of contigs | 1933851 | 17897 | 67266 | 26988 |
| Smallest contig | 64 | 231 | 200 | 200 |
| Largest Contig | 94914 | 162047 | 16167 | 16167 |
| Number of bases | 631125096 | 194706849 | 29298709 | 13126985 |
| Mean contig length | 237.49186 | 10879.30094 | 435.56 | 486.40 |
| n_under_200 | 1679726 | 0 | 0 | 0 |
| Number of contigs over 1k | 107943 | 17846 | 1456 | 832 |
| Number of contigs over 10k | 5686 | 6597 | 3 | 2 |
| n90 | 809 | 5106 | 268 | 301 |
| n70 | 2774 | 9988 | 363 | 415 |
| n50 | 6790 | 15425 | 458 | 530 |
| n30 | 23809 | 25478 | 592 | 668 |
| n10 | 94914 | 49505 | 908 | 975 |
| gc% | 0.37927 | 0.38219 | 0.44 | 0.45 |
| Number of bases that are N | 361257 | 439633 | 0 | 0 |
| Proportion of bases that are N | 0.00057 | 0.00226 | 0 | 0 |

**S4 Table.** Summary of SNPs used in the analysis of linkage disequilibrium. Only SNPs that could be mapped to the genome of the close relative, *I. nil*, were used in analyses. r^2^ values were determined using all individuals regardless of population or resistance level.

| Chromosome | SNP Number | $r^2$ mean | SD | $r^2$ 75% | $r^2$ within 10kb |
| --- | --- | --- | --- | --- | --- |
| 1 | 1488 | 0.036 | 0.085 | 0.033 | 0.071±0.02 |
| 6 | 3191 | 0.032 | 0.077 | 0.031 | 0.038±0.001 |
| 10 | 1779 | 0.033 | 0.074 | 0.033 | 0.057±0.009 |
| 13 | 2011 | 0.034 | 0.077 | 0.034 | 0.039±0.004 |
| 15 | 1189 | 0.035 | 0.087 | 0.032 | 0.078±0.02 |

**S5 Table.**
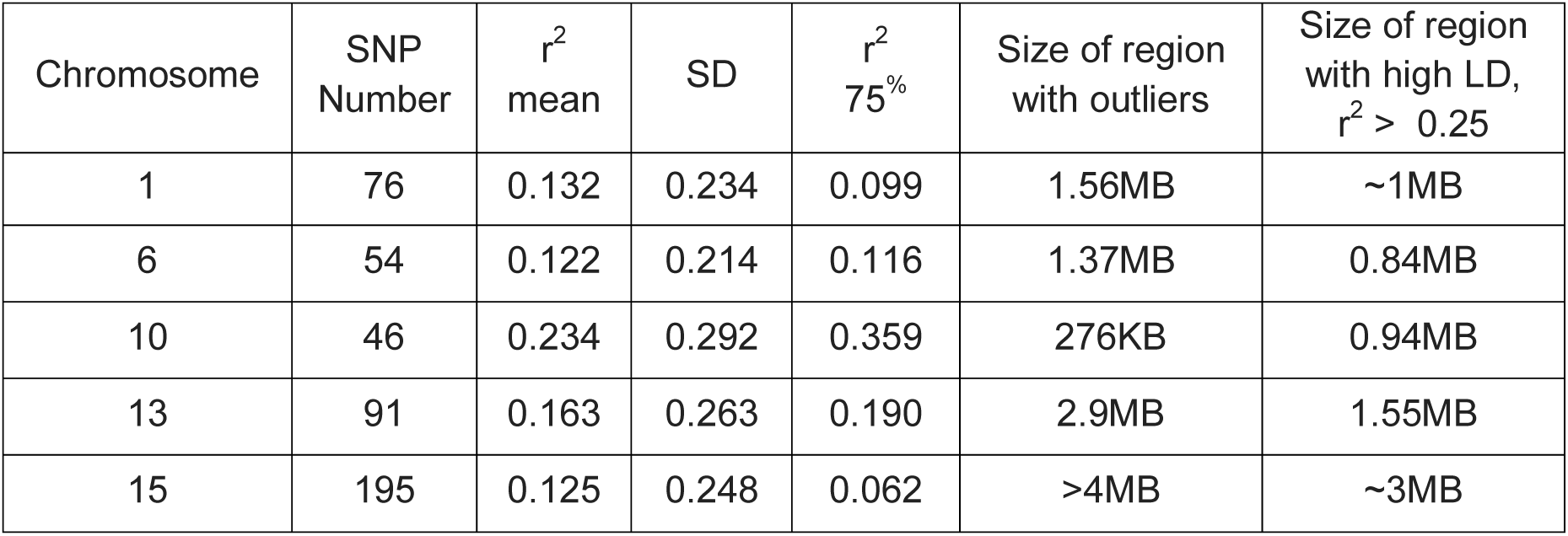
Summary of SNPs used in the analysis of linkage disequilibrium in the regions enriched for outliers, per chromosome, as identified by bayenv2 or Bayescan. Only SNPs that could be mapped to the genome of the close relative, *I. nil*, were used in analyses. r^2^ values were determined using all individuals regardless of population or resistance level.

| Chromosome | SNP Number | $r^2$ mean | SD | $r^2$ 75% | Size of region with outliers | Size of region with high LD, $r^2 > 0.25$ |
| --- | --- | --- | --- | --- | --- | --- |
| 1 | 76 | 0.132 | 0.234 | 0.099 | 1.56MB | ~1MB |
| 6 | 54 | 0.122 | 0.214 | 0.116 | 1.37MB | 0.84MB |
| 10 | 46 | 0.234 | 0.292 | 0.359 | 276KB | 0.94MB |
| 13 | 91 | 0.163 | 0.263 | 0.190 | 2.9MB | 1.55MB |
| 15 | 195 | 0.125 | 0.248 | 0.062 | >4MB | ~3MB |

**S6 Table.**
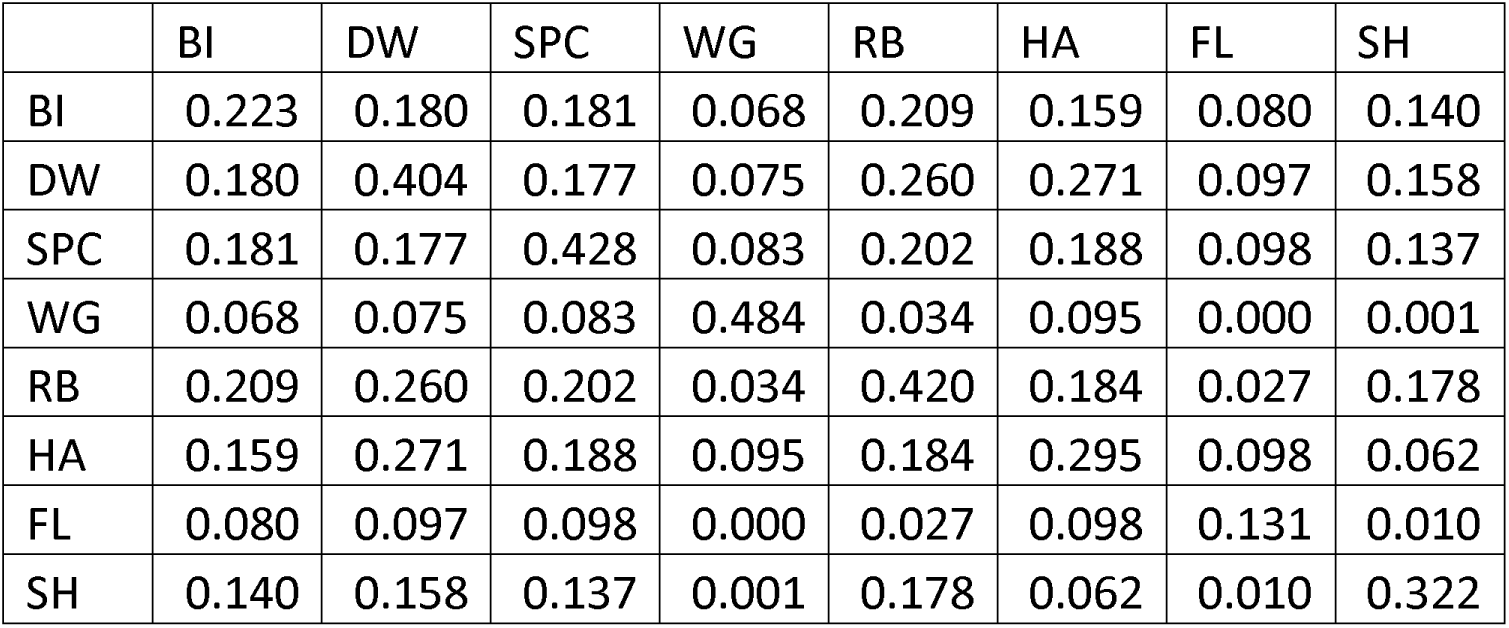
Neutral F matrix from scaffolds on chromosomes 3, 7, and 14 (61 scaffolds total). BI, DW, SPC, and WG are the high resistance populations.

|  | BI | DW | SPC | WG | RB | HA | FL | SH |
| --- | --- | --- | --- | --- | --- | --- | --- | --- |
| BI | 0.223 | 0.180 | 0.181 | 0.068 | 0.209 | 0.159 | 0.080 | 0.140 |
| DW | 0.180 | 0.404 | 0.177 | 0.075 | 0.260 | 0.271 | 0.097 | 0.158 |
| SPC | 0.181 | 0.177 | 0.428 | 0.083 | 0.202 | 0.188 | 0.098 | 0.137 |
| WG | 0.068 | 0.075 | 0.083 | 0.484 | 0.034 | 0.095 | 0.000 | 0.001 |
| RB | 0.209 | 0.260 | 0.202 | 0.034 | 0.420 | 0.184 | 0.027 | 0.178 |
| HA | 0.159 | 0.271 | 0.188 | 0.095 | 0.184 | 0.295 | 0.098 | 0.062 |
| FL | 0.080 | 0.097 | 0.098 | 0.000 | 0.027 | 0.098 | 0.131 | 0.010 |
| SH | 0.140 | 0.158 | 0.137 | 0.001 | 0.178 | 0.062 | 0.010 | 0.322 |

**S7 Table.** Parameter spaces for composite-likelihood calculations for the standing variation (s, t, g) and migration (s, m, source population) model simulations.

|  |  |
| --- | --- |
| Position of selected site | 37189, 37198, 37224, 37246, 37258, 37267, 37271, 37273, 37282, 37283, 37285, 37288, 37303, 37305, 37342, 37355, 37357, 37360, 37362, 37366, 37376, 37408, 140310, 140466, 140544, 140552, 140565, 140571, 140605, 140627 |
| s | 0.001, 0.002, 0.003, 0.004, 0.005, 0.006, 0.007, 0.008, 0.009, 0.01, 0.011, 0.014, 0.016, 0.019, 0.021, 0.024, 0.026, 0.029, 0.032, 0.034, 0.037, 0.039, 0.042, 0.045, 0.047, 0.05, 0.052, 0.055, 0.057, 0.06, 0.08, 0.1, 0.15, 0.2, 0.25, 0.3, 0.35, 0.4, 0.45, 0.5, 0.55, 0.6, 0.65, 0.7, 0.75, 0.8, 0.85, 0.9, 0.95, 1 |
| t | 5, 10, 81, 151, 222, 293, 364, 434, 505, 576, 646, 717, 788, 859, 929, 1000, 1500, 1607, 1714, 1821, 1929, 2036, 2143, 2250, 2357, 2464, 2571, 2679, 2786, 2893, 3000 |
| g | $10^{-10}$, $10^{-9}$, $10^{-8}$, $10^{-7}$, $10^{-6}$, $10^{-5}$, $10^{-4}$, $10^{-3}$, $10^{-2}$ |
| m | $10^{-5}$, $10^{-4}$, $5^{-4}$, 0.001, 0.005, 0.01, 0.1, 0.2, 0.3, 0.4, 0.5, 0.6, 0.7, 0.8, 0.9, 1 |
| source population | SPC and WG |

**S1 Dataset:** Tables include annotations of outlier RADseq loci, annotations of probe sequences used for target capture probes, annotation of outlier contigs from resequencing, a list of *I. nil* genes within the 5 outlier enriched regions. https://docs.google.com/spreadsheets/d/1I59RoHSTc4ktXMOuZQuN5KNxMQ0Lprozqma8Cf3gBmA/edit?usp=sharing

## Acknowledgements

We thank A. Kuester for seed collection, A. Wilson for assistance with lab work, M.C. Hwang for assistance with initial genome assembly, E. Schold for assistance with EPSPS sequencing, and S.D. Smith, G. Coop, M. Hahn, and J. Ross-Ibarra for providing comments on an earlier version of this work. MVE, S-MC and RSB were supported by USDA XXX, and KL was supported by NIH XXX.

## Author contributions

Conceptualization: ME Van Etten, RS Baucom

Data curation: ME Van Etten

Formal analysis: ME Van Etten, RS Baucom, KM Lee

Funding acquisition: RS Baucom, S-M Chang

Investigation: ME Van Etten, RS Baucom, KM Lee

Methodology: ME Van Etten, RS Baucom, KM Lee

Supervision: RS Baucom

Visualization: ME Van Etten, RS Baucom

Writing --original draft: ME Van Etten, KM Lee, S-M Chang, RS Baucom

